# Human Accelerated Regions regulate gene networks implicated in apical-to-basal neural progenitor fate transitions

**DOI:** 10.1101/2024.06.30.601407

**Authors:** Mark Alan Noble, Yu Ji, Kristina M. Yim, Je Won Yang, Matheo Morales, Reem Abu-Shamma, Atreyo Pal, Ryan Poulsen, Marybeth Baumgartner, James P. Noonan

**Affiliations:** Department of Genetics, Yale School of Medicine, New Haven, CT 06510, USA; Department of Neuroscience, Yale School of Medicine, New Haven, CT 06510, USA; Wu Tsai Institute, Yale University, New Haven, CT 06510, USA

## Abstract

The evolution of the human cerebral cortex involved modifications in the composition and proliferative potential of the neural stem cell (NSC) niche during brain development. Human Accelerated Regions (HARs) exhibit a significant excess of human-specific sequence changes and have been implicated in human brain evolution. Multiple studies support that HARs include neurodevelopmental enhancers with novel activities in humans, but their biological functions in NSCs have not been empirically assessed at scale. Here we conducted a direct-capture Perturb-seq screen repressing 180 neurodevelopmentally active HARs in human iPSC-derived NSCs with single-cell transcriptional readout. After profiling >188,000 NSCs, we identified a set of HAR perturbations with convergent transcriptional effects on gene networks involved in NSC apicobasal polarity, a cellular process whose precise regulation is critical to the developmental emergence of basal radial glia (bRG), a progenitor population that is expanded in humans. Across multiple HAR perturbations, we found convergent dysregulation of specific apicobasal polarity and adherens junction regulators, including *PARD3, ABI2, SETD2*, and *PCM1*. We found that the repression of one candidate from the screen, HAR181, as well as its target gene *CADM1*, disrupted apical PARD3 localization and NSC rosette formation. Our findings reveal interconnected roles for HARs in NSC biology and cortical development and link specific HARs to processes implicated in human cortical expansion.

## Introduction

The evolutionary expansion and elaboration of the human cerebral cortex is in part the product of modifications to neural stem cell (NSC) fate specification, migration, proliferation, and the timing of neural differentiation in development^1–3^. Genetic changes that altered developmental enhancer activity have been hypothesized to drive many of these traits^4–7^. Human Accelerated Regions (HARs) are genomic regions that are highly constrained across multiple species, but which show a significant excess of human-specific substitutions (**Fig. 1A**)^8–11^. This evolutionary acceleration suggests HARs encode potential human-specific functions, and several lines of evidence support that many HARs act as neurodevelopmental enhancers exhibiting human-specific changes in activity^12–18^. Studies of three-dimensional chromatin contacts between HARs and genes support that HARs regulate genes with neurodevelopmental functions^19–21^. Massively parallel reporter assays (MPRAs) have identified HARs with differential activity compared to their chimpanzee orthologs in a variety of cell types, including in human NSCs (hNSCs)^15,16,22^. A recent study using CRISPR perturbation to disrupt thousands of enhancers in hNSCs identified 15 HARs which were required for hNSC self-renewal^18^.

**Figure 1.**
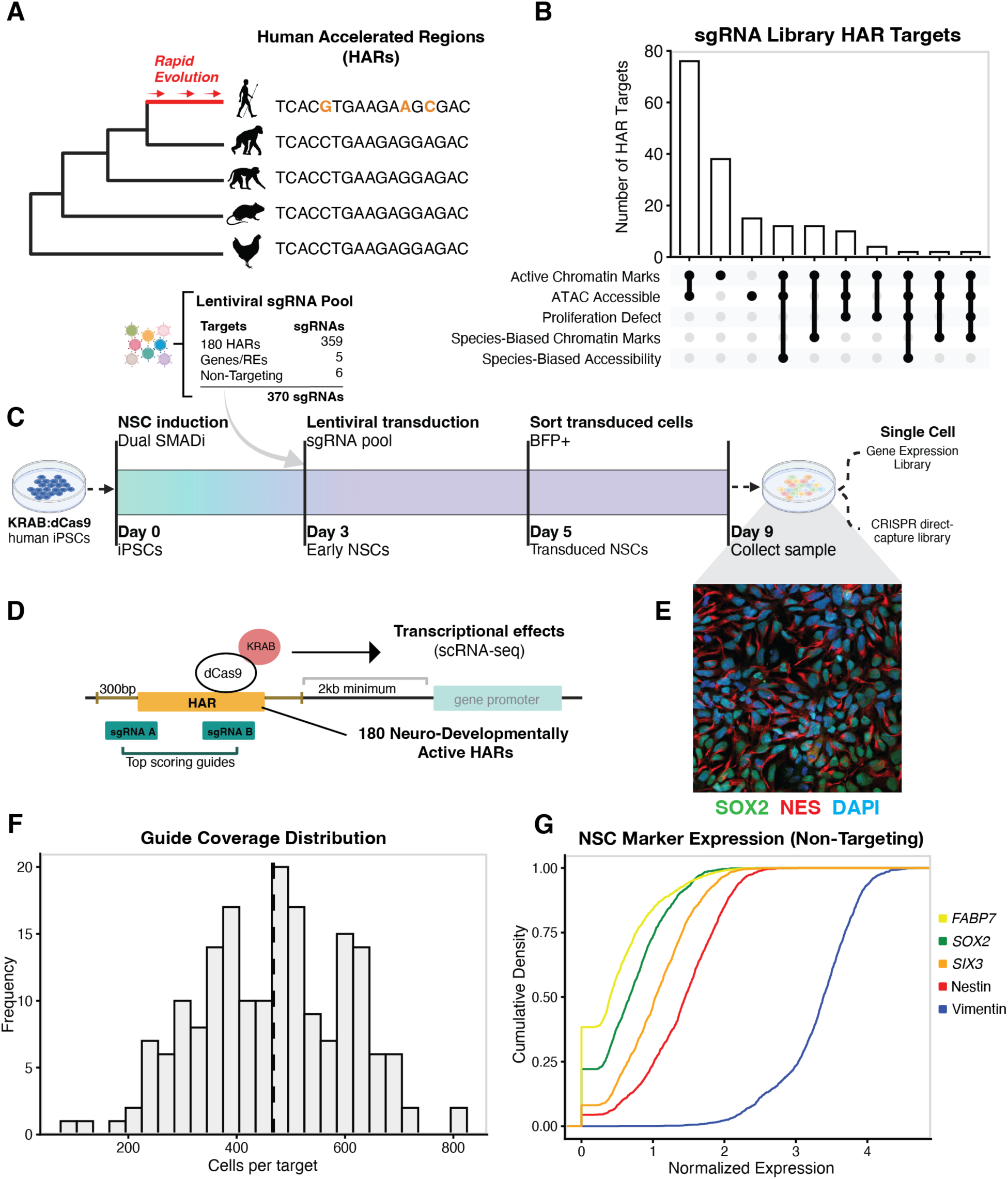
A direct-capture Perturb-seq screen targeting 180 HARs in human neural stem cells. **A.** Schematic illustrating how Human Accelerated Regions are defined. **B.** Upset graph showing the number of HAR targets represented in the Perturb-seq sgRNA library that intersect each of the annotations considered as evidence of neurodevelopmental or species-biased enhancer activity. **C.** Schematic of the direct-capture Perturb-seq experiment; see Methods for details. **D.** Schematic illustrating how sgRNAs were targeted to each HAR. Each was targeted by two high-scoring sgRNAs (Methods) within 300bp of the HAR. Targeted regions were located at least 2kb from any nearby promoter, if present. **E.** Expression of the NSC markers SOX2 and NES in hNSCs carrying KRAB:dCas9. **F.** The distribution of the number of high-quality cells in which an sgRNA could be assigned for each target. After filtering for cells bearing a single sgRNA, targets were represented by an average of ∼500 cells. **G.** Cumulative density plot showing the expression distribution of the indicated NSC marker genes in NTC-sgRNA bearing cells in the Perturb-seq screen. Expression values are normalized to the total number of UMI counts for each cell, multiplied by a scaling factor (1000), then log transformed. Schematics were generated in part using BioRender.com.

As HAR targets have been shown to be enriched in particular cellular pathways and for ontologies associated with multiple neurodevelopmental processes, it has been hypothesized that some HARs might have convergent regulatory roles^5,17,21^. However, the specific biological processes that HARs regulate are largely unknown and experimental data supporting a coordinated role for HARs in cortical development are limited. To address these questions, we carried out a pooled CRISPR screen with single-cell RNA-seq readout and direct capture of sgRNAs, an approach termed direct-capture Perturb-seq^23^, to repress 180 HARs in human iPSC-derived NSCs. While this method has primarily been used to directly perturb genes, previous studies have quantified transcriptional effects by perturbing enhancers^24,25^. We reasoned that silencing HARs at scale in a single, internally-controlled experiment would allow us to characterize their downstream, and potentially convergent, regulatory functions.

We profiled over 188,000 cells and then modeled the effects of HAR perturbations on functionally-annotated co-expression modules using multiple linear regression. We identified numerous HARs with significant effects on gene modules regulating diverse cellular processes. We also found that a subset of HAR perturbations downregulated gene networks associated with the establishment and maintenance of the NSC apical domain and cell polarity, cellular functions with specific roles in modulating the transition from apical to basal radial glia, a process which is critical to cortical expansion and has been implicated in human evolution^26^. Further, we identified specific genes on which the transcriptional changes resulting from multiple HAR perturbations converged, including *PARD3*, *ABI2*, *PCM1*, and *SETD2*, important regulators of NSC cell adhesion, cytoskeleton, and NSC fate decisions. Repression of one HAR whose perturbation resulted in strong transcriptional downregulation of polarity-associated genes, HAR181, as well as its target gene *CADM1*, resulted in disruption of NSC rosette formation and mislocalization of *PARD3* in hNSCs. Together, our findings suggest that multiple HARs contribute to the regulation of NSC apicobasal polarity, a feature implicated in the evolution and expansion of the cortical proliferative niche. Our results thus further implicate HARs in neurodevelopmental processes involved in human brain evolution.

## Results

### Using direct-capture Perturb-seq to investigate HAR neurodevelopmental functions

Previous direct-capture Perturb-seq studies have successfully identified transcriptional effects due to gene and enhancer perturbations in multiple biological contexts^27–30^. However, prior work also indicates that statistical power to detect differential gene expression relies on adequate representation of target sgRNAs in cells as well as sufficient sequencing depth, considerations that limit the number of loci that can be effectively targeted in a single screen^31^. Therefore, we prioritized 180 HARs which had strong evidence for neurodevelopmental regulatory activity, such as chromatin accessibility or marking by the active enhancer-associated histone modification H3K27ac, in published datasets from human fetal brain (RoadMap consortium) and organoid studies (**Fig. S1B**; Methods). We further prioritized HARs which showed differences in chromatin accessibility in human compared to chimpanzee cerebral organoids, HARs which were required for hNSC proliferation, and HARs that overlapped Human Gain Enhancers, a class of putative enhancers defined by increased H3K27ac or H3K4me2 marking in human fetal cortex compared to rhesus macaque and mouse (Methods)^18,32,33^. Of the HARs targeted in the final sgRNA library, 104 were both active and accessible during brain development, 20 were associated with NSC proliferation defects, and 22 showed evidence of a species-bias in accessibility or activity (**Fig. 1B**).

We generated an sgRNA lentiviral library to target each HAR with high specificity (**Fig. S1A**; Methods). We stringently filtered sgRNAs for high scores for on-target specificity and with no potential off-target sites that had fewer than 3 mismatches (**Fig. S1A**, **S1C-D**). For each target, 2 sgRNAs were selected (except for one HAR, *HACNS255*/HAR195, for which only 1 unique sgRNA could be designed; **Fig. 1D**). This yielded a library of 370 sgRNAs targeting 180 HARs, as well as 6 scrambled non-targeting controls (NTCs) and 5 sgRNAs targeting previously validated positive control loci^18,34^. All sgRNA sequences and their genomic locations (GRCh38 coordinates) are provided in **Table S1**. We then generated polyclonal cell lines expressing KRAB:dCas9 from human iPSC lines via low-MOI lentiviral transduction and Blasticidin selection (Methods). These were then differentiated into NSCs using dual SMAD inhibition. We verified expression of KRAB:dCas9 in both iPSCs and differentiated NSCs, as well as stable expression of NSC markers (e.g. *PAX6*, *SOX2*, *NES*) via RT-qPCR, and NSC morphology and marker expression (SOX2 and NES) via immunofluorescence microscopy^35,36^ (**Fig. 1E, Fig. S2A-B**). We further validated knockdown competency in the KRAB:dCas9 cell line by introducing an sgRNA targeting the promoter of *GRN*, which resulted in a 42-84% reduction in *GRN* expression compared to control^18,34^ (**Fig. S2C-D**). Perturb-seq was then performed as shown in **Fig. 1C**. We profiled 188,969 cells across 3 batches. For our analyses, we only considered 87,896 cells with a single sgRNA assignment and which passed quality control filters (**Fig. S4A-B**; Methods). The distribution of sgRNA assignment rates was consistent across batches (**Fig. S4C-D**), and sgRNAs were highly represented, with an average of ∼450 cells/sgRNA (**Fig. 1F**). We observed consistent expression of canonical NSC markers in cells with NTC sgRNAs^35–38^, supporting that lentiviral transduction and subsequent steps did not alter NSC identity (**Fig. 1G**).

### Weighted gene correlation analysis identifies gene co-expression modules in perturbed NSCs

We first sought to identify the transcriptional effects of HAR perturbation by identifying differentially expressed genes (DEGs) in perturbed versus NTC sgRNA-bearing cells using Wilcoxon rank sum tests, then intersecting these DEGs with HAR gene targets previously identified in hNSCs^21^. We identified 23 perturbations that had negative effects (log2 fold change < -0.1) on the expression of their predicted target genes (**Fig. S5**). These included 3 separate perturbations (2xHAR.446, *HACNS927*, *HACNS839*) which resulted in downregulation of their common target, Neurotrimin (*NTM*), a regulator of cell adhesion and neurite outgrowth. Positive control loci similarly showed negative fold changes in gene expression (**Fig. S6A-E**). However, for some positive controls, larger effect sizes were seen for genes which were not directly targeted (**Fig. S6F-J**), indicating we had a greater ability to detect downstream transcriptional effects, potentially for genes with higher expression levels than the primary target genes. This is consistent with previous Perturb-seq studies, which have faced limitations in identifying significant, biologically relevant effects from pseudo-bulk differential expression approaches alone, instead utilizing other approaches, including methods intended to isolate the signal resulting from the perturbation of known gene targets^39,40^. Other studies have also broadened their analyses to ontological gene groups or gene co-expression modules to detect global transcriptional effects resulting from perturbation^41,42^.

Therefore, to further understand the functional effects of HAR perturbation, we employed weighted gene correlation network analysis (WGCNA), as described in a previous study^41^, to define modules of co-expressed genes in the Perturb-seq expression data (**Fig. 2A**)^43^. In this method, each expressed gene is assigned to a single module consisting of genes with highly correlated gene expression, or to no module (M0). Such gene modules have previously been shown to reveal sets of co-regulated genes expressed in specific cell types or that operate in specific pathways that can be identified using Gene Ontology (GO) analysis^44,45^. Using the hdWGCNA package^43^, we identified 16 co-expression modules and performed Gene Ontology enrichment analysis to determine biological functions associated with each (**Fig. 2B**; Methods). We identified modules enriched for genes involved in DNA synthesis (M7), translation (M8 and M13), and mitochondrial function (M4 and M15). Additionally, we identified modules enriched for terms associated with processes known to play important roles in NSC fate decisions and progenitor subtype specification during neurodevelopment, such as apicobasal polarity and Rho-GTPase regulation (M3 and M5), stress and Wnt signaling (M9), and cell migration (M2 and M13). A full list of module genes is available in **Table S4**, and GO term enrichments are available in **Table S5**.

**Figure 2.**
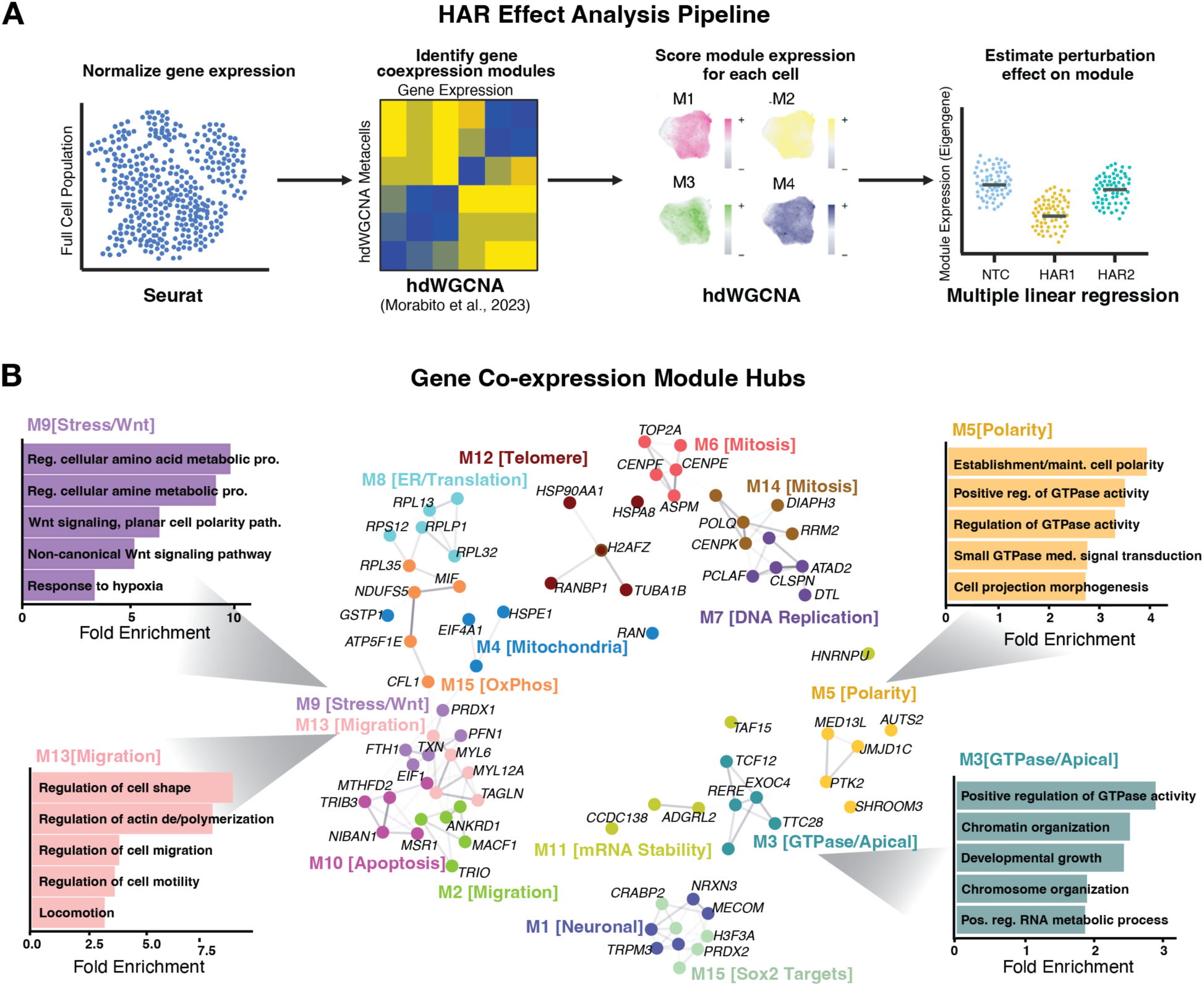
Weighted gene co-expression network analysis identifies 16 functional NSC gene modules. **A.** The analytical pipeline used to quantify HAR perturbation effects on gene modules identified from the Perturb-seq assay. hdWGCNA was applied to the population of 87,896 high quality cells to identify gene networks. Module expression (defined based on the module eigengene; see Results and Methods) was scored for each cell, and multiple linear regression was performed to assess the overall effects of HAR perturbation of each module’s expression. This schematic was generated using BioRender.com. **B.** The top 5 module hub genes, defined as the 5 genes that are most highly correlated with their respective module’s expression. Each point represents a hub gene, with edges indicating expression correlations among hub genes. Hubs are colored by module membership. Gene Ontology Biological Process enrichments for highly significant terms within key modules are shown at each side of the figure.

### HAR perturbations alter gene co-expression network expression

We next asked what the effects of HAR perturbations were on module expression, defined as module eigengene values (MEs), which are defined as the first principal component of the expression matrix of genes assigned to a given module. We fit an additive multiple linear regression (MLR) model for the expression of each module in response to the following covariates: sgRNA assignment, batch, and number of genes detected per cell (Methods; **Fig. 2A**). Using this model, we estimated effect sizes and significance (false discovery rate adjusted P-values; FDR) for each HAR perturbation on each module’s expression compared to the module’s expression in cells bearing NTC sgRNAs, while accounting for batch and cell library size. Effect sizes are thus defined as the estimated shifts in module expression with respect to cells bearing NTC sgRNAs.

The distributions of HAR perturbation effect sizes for each module are shown in **Figure 3A**. In all, 12 of the 16 modules were significantly affected by at least one HAR perturbation (FDR < 0.05). The number of perturbations significantly affecting each module are summarized in **Table S8**. The estimated effect sizes for the batch covariate were also significant, indicating our model accounted for variation in module expression across batches. We found no correlation between the number of genes within each module and the number of significant effects on that module’s expression resulting from HAR perturbations (Pearson correlation, R = -0.041, p = 0.9) indicating the rate of discovery was not biased by module size (**Fig. S8C**). In total, 27 HAR perturbations resulted in significant disruption of one or more modules (FDR < 0.05; **Fig. 3B**). Of these, many HAR perturbations yielded similar effects, with several perturbations resulting in significant downregulation of modules M3 and M5, both involved in apicobasal polarity, and upregulation of modules such as M9, M13, and M15, which are enriched for genes involved in stress, migration, and metabolism, respectively (**Fig. 3C**).

**Figure 3.**
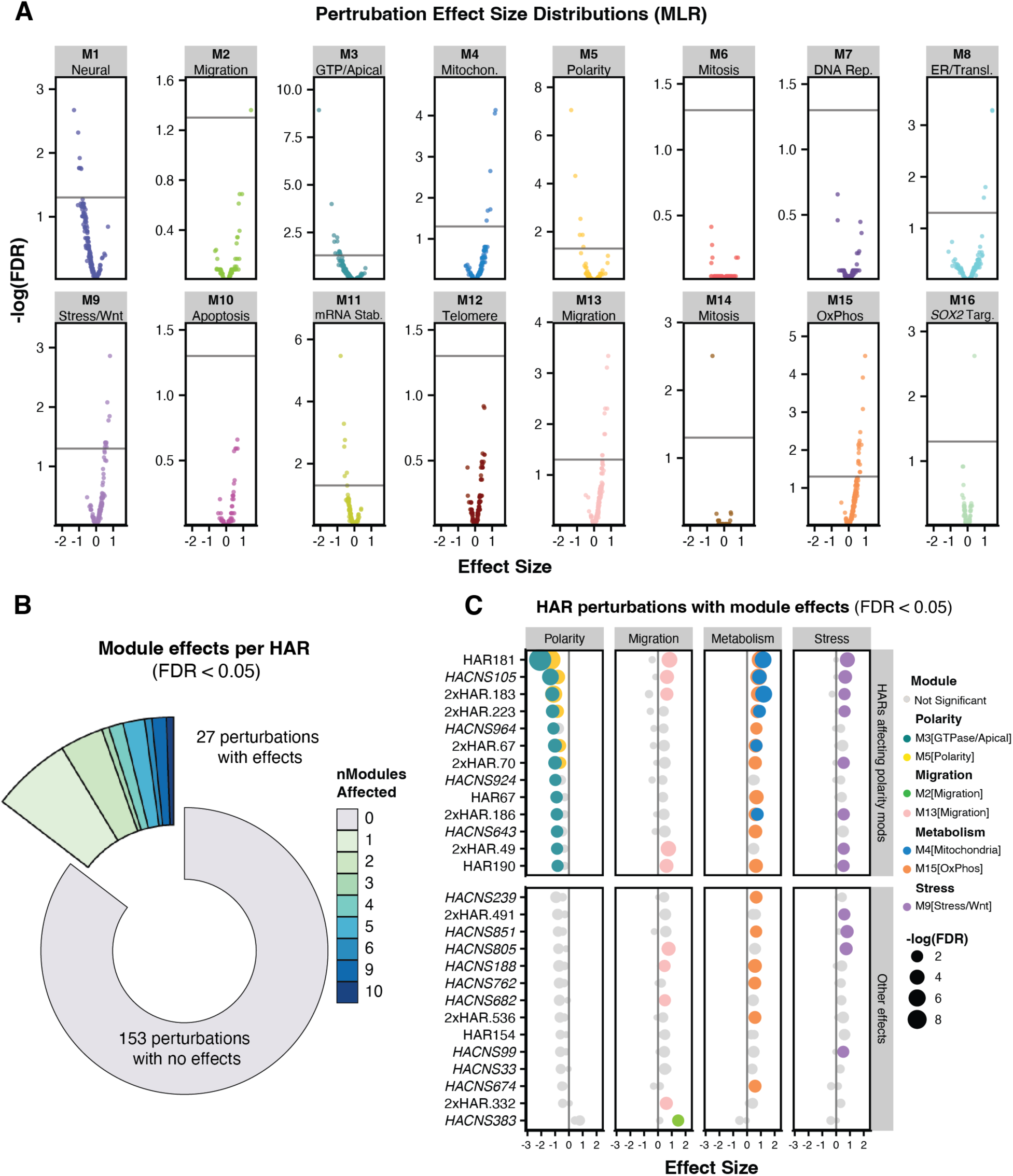
A subset of HAR perturbations affect gene module expression. **A.** Effect sizes for HAR perturbations on gene module expression were estimated by multiple linear regression (MLR). The distribution of effect sizes in reference to NTC sgRNA-bearing cells, and associated FDRs, for each module’s expression from the MLR model are shown. Each point represents the overall effect of each HAR perturbation on that module. Points are colored by module membership as shown in Figure 2B. The horizontal line on each plot indicates an FDR significance threshold of 0.05. **B**. Donut plot showing the distribution of the number of significant effects (FDR < 0.05) for each HAR perturbation on module expression from the MLR model. **C**. A subset of HAR perturbations with significant effects. All perturbations with significant effects are shown in Figure S8. HARs are presented on the Y-axis. The upper panel shows HAR perturbations that significantly affect modules associated with polarity, while the lower panel shows perturbations that affect modules associated with other functions. Module expression effect sizes from the MLR model are plotted on the X-axis, and significance (-log10 FDR) is represented by point size. Module membership is represented by point color, and modules are categorized into general ontologies as in Figure 2B. Modules not showing significant effects (FDR < 0.05) are plotted in grey.

Of the 13 perturbations affecting the GTPase/apical module M3, 6 also showed significant negative effect sizes for the polarity-associated module M5 (**Fig. 3C**). These included perturbations of HAR181, *HACNS105*, 2xHAR.223, and 2xHAR.183, all of which also resulted in increased expression of modules associated with stress (M9), migration (M13), and oxidative phosphorylation (M15). Additionally, 7 perturbations downregulated module M1, which included genes involved in neural lineage specification (**Fig. S8A**). Six perturbations downregulated module M11, enriched for genes regulating mRNA stability and transport. Only one perturbation had any effects on modules associated with the cell cycle (2xHAR.183, with a negative effect size for mitosis-associated module M14), and no perturbations affected modules associated with DNA replication, telomere maintenance, or apoptosis. These results support that HAR perturbations result in broad effects on gene network expression related to a wide range of cellular processes, with some perturbations resulting in the dysregulation of multiple gene modules. HAR perturbations showed convergent patterns in module dysregulation, with consistent downregulation of polarity-associated modules, and upregulation of modules related to migration, metabolism, and stress.

### HAR perturbations disrupt the expression of genes involved in NSC apicobasal polarity

We found that multiple HAR perturbations downregulated modules associated with apicobasal polarity. In early cortical development the proliferating neuroepithelium and apical radial glia (aRG) are affixed by adhesions to the ventricular surface via apical adherens junctions, which physically limits the number of possible aRG divisions^2,46^ (**Fig. 4A**). These adherens junctions are established and maintained by apicobasal polarity genes including *PARD3* and *AFDN*^47–50^. As development progresses, asymmetric divisions give rise to basal radial glia (bRG), which do not reestablish apical domains, while migratory pathways are activated, releasing progenitors from the ventricular surface^49,50^. In this manner, the process of progenitor delamination expands the proliferative niche of the developing cortex. Such bRG cells are more abundant in the developing human cortex than in other primates, and this is hypothesized to contribute to cortical expansion in evolution^49–51^. Prior studies have demonstrated that cultured NSCs exhibit features of apicobasal polarity in culture, ^52,53^ and we observed this in our culture system, as NSCs form rosette structures with inward-facing apical junctions marked with *PARD3* by day 9 of NSC induction (**Fig. 4B**). Thus, aspects of the gene networks and cellular systems maintaining apicobasal polarity in NSCs are recapitulated in our monolayer culture system.

**Figure 4.**
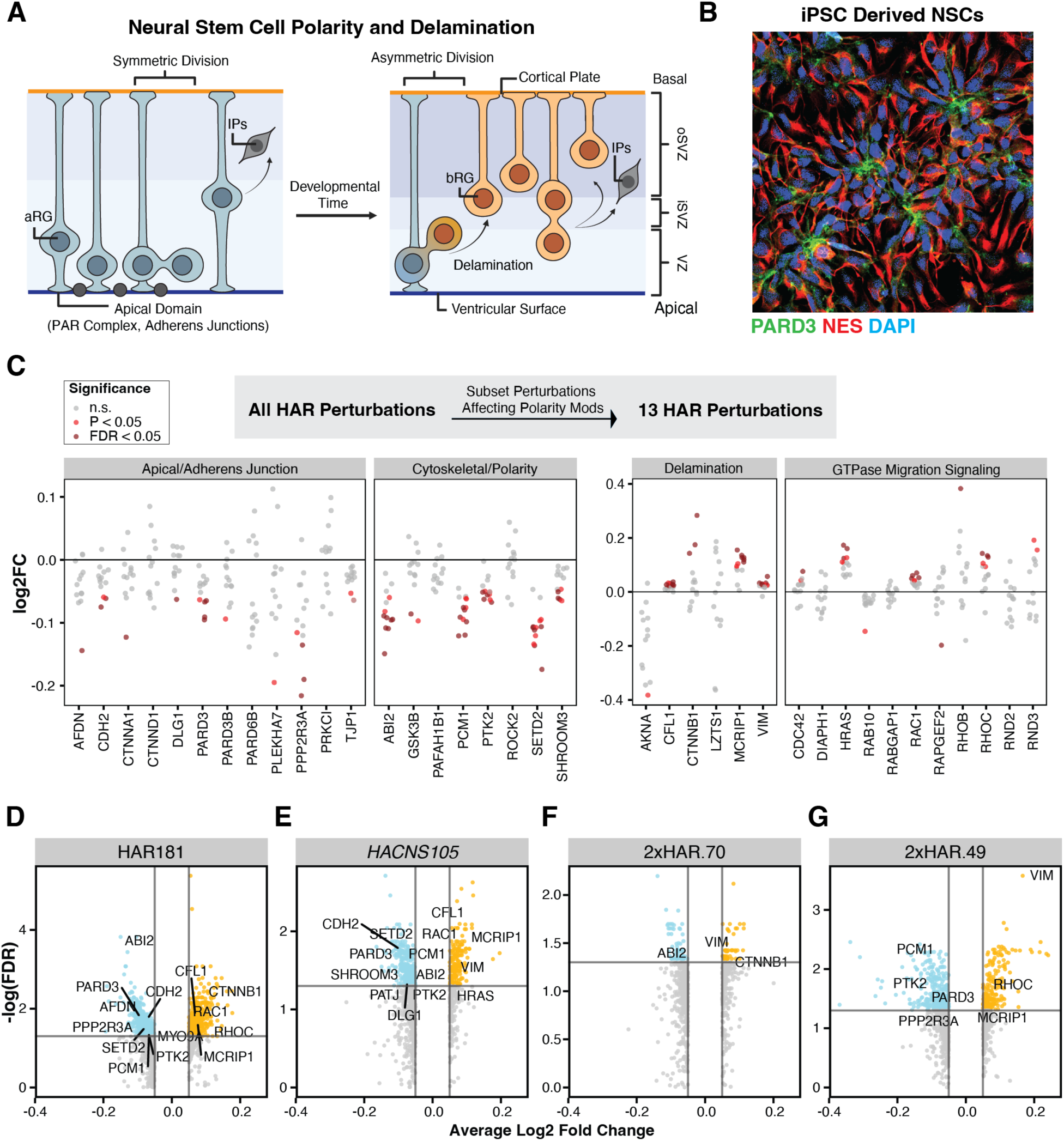
A subset of HAR perturbations affect regulators of NSC polarity and migration. **A.** Schematic summarizing NSC delamination and bRG specification in cortical development. *Left*. Cortical NSCs exhibit apicobasal polarity as aRG early in development *in vivo*. Cells express genes such as those in the PAR complex which establish and maintain the apical domain, which anchors the cells to the ventricular surface. *Right.* The proliferative niche of the developing cortex is expanded as aRGs undergo asymmetric divisions to yield cells lacking apical domains, the bRG, which delaminate and are free to migrate basally. **B**. Immunofluorescence microscopy of iPSC-derived (polyclonal KRAB:dCas9 line) NSCs from day 9 post induction exhibiting rosettes that show asymmetric localization of PARD3 (green) towards the interior cell junctions. NES is shown in red and DAPI nuclear staining is shown in blue. **C.** Differential gene expression (Log2 fold change compared to NTC cells) from 13 HAR perturbations in the Perturb-seq screen affecting polarity-associated modules. Genes associated with the apical domain, polarity and cytoskeleton regulation are shown in the left panels, and genes associated with delamination and migration are shown in the right panels. For each panel, gene names are plotted on the X-axis and fold changes are plotted on the Y-axis. Fold changes for all 13 perturbations are shown for each gene. Significant values from Wilcoxon Rank Sum tests are indicated in red. **D-G.** Volcano plots of log2 fold changes plotted against -log10 FDR for 4 HAR perturbations affecting polarity modules. Significantly downregulated genes are colored in teal, and significantly upregulated genes are colored in orange. Notable DEGs associated with NSC polarity or migration processes are labeled.

Given our finding that a subset of HAR perturbations disrupted gene modules involved in apicobasal polarity, we reevaluated the same sets of DEGs we previously calculated for the Perturb-seq screen, focusing on genes established to be involved in the establishment of the apical domain and adherens junctions, and the regulation of the cytoskeleton and NSC polarity^48,54–60^(**Fig. 4C**). We found that, of 13 perturbations which caused downregulation in modules associated with apicobasal polarity, all resulted in significant downregulation of genes important for the establishment and maintenance of the aRG apical domain, including members of the PAR complex (*PARD3*, *PARD3B*), ZO-1 (*TJP1*), Afadin (*AFDN*), and E-Cadherin (*CDH2*). Similarly, several cytoskeletal modifiers such as *ABI2* and *PCM1*, genes involved in polarity maintenance and mitotic spindle orientation^61,62^, processes which are crucial for the regulation of the aRG to bRG transition, were also downregulated. In addition, we detected significant upregulation of genes involved in NSC delamination and migration (**Fig. 4C**). Delamination and migration-associated genes including Vimentin (*VIM*) and *MCRIP*, were upregulated. Multiple Rho GTPases and related genes, which coordinate NSC shape and migration processes^54,63–65^, were also upregulated, including *CDC42*, *RHOB*, *RHOC*, and *RAC1*. Across the differential expression profiles for these perturbations, we observed concurrent downregulation of apical-domain and polarity associated genes coupled with the upregulation of migration-associated genes (**Fig. 4D-F**).

We further investigated the effects of HAR perturbation on apicobasal polarity gene expression by using CRISPRi to specifically silence HAR181 in a KRAB:dCas9-expressing NSCs derived from an independent iPSC cell line. Perturbation of HAR181 yielded the strongest downregulation of the GTPase/apical-associated module M3 and the polarity-associated module M5 in our screen. Using RT-qPCR to quantify gene expression, we found that repressing HAR181 resulted in significant downregulation of its gene target, *CADM1*, compared to cells transfected with NTC sgRNAs (**Fig. S9**). The polarity-related genes *PARD3, SETD2* and, *AFDN* were also significantly downregulated, as we observed in the screen. The downregulation of *PARD3* was also recapitulated by CRISPRi silencing of *CADM1*, supporting that this phenotype is the result of HAR181-mediated regulation of *CADM1* expression. We also evaluated another downregulated DEG identified in our screen, *ABI2*, but we did not observe significant downregulation of this target due to HAR181 or *CADM1* repression. Collectively, our results identify a subset of HAR perturbations that converge on the downregulation of apical-domain-specifying genes critical to aRG cell identity. Furthermore, the perturbation of HAR181 support provides orthogonal support for the observed transcriptional phenotypes from our screen.

### Overlapping gene-level effects of HAR perturbations affecting polarity-associated modules

Given their convergent effects on module expression, we further investigated whether the 13 HAR perturbations which resulted in the downregulation of polarity-associated modules (**Fig. 3C**) affected the expression of common sets of genes. We assessed the DEGs resulting from these 13 HAR perturbation expression profiles and tabulated the number of HAR perturbations which caused significant down- or upregulation of each gene. We found that some genes were affected by up to 9 of the 13 perturbations shown in **Figure 4C**, indicating significant overlap in the downstream effects of independent perturbations (**Fig. S9**). Fifty-one genes in the GTPase/apical-associated module M3 were downregulated by >3 perturbations: 28 were downregulated by 3 perturbations, 15 were downregulated by 4 perturbations, 5 were downregulated by 5 perturbations, and 3 genes were downregulated by 6 perturbations (**Fig. 5A**). Genes without a module assignment (M0) and neural identity related genes (M1) were also affected by multiple perturbations, indicating broader downregulation of genes outside of polarity networks alone. Across all modules, 9 genes in total were downregulated by 6 or more perturbations (**Fig. 5C**). These included *ABI2*, a member of the polarity-associated module M5 and a regulator of Arp2/3 actin polymerization and cell migration in the developing cortex^61^; *SETD2*, a chromatin and microtubule modifier required for neural cell shape^66–70^; and *PCM1*, a regulator of mitotic spindle orientation which is required for RG proliferation and is involved in bRG specification^62,70,71^. The largest number of genes upregulated by 3 or more perturbations belonged to the mitochondria-associated module M4 and the migration-associated module M13, suggesting a convergent upregulation of metabolic and migratory pathways due to perturbation (**Fig. 5B**). Fourteen genes were upregulated by 6 or more perturbations, including genes such as *PRDX1*, the antioxidant stress gene periodoxin, and *GPX4*, an antioxidant which responds to lipid peroxidation^72,73^ (**Fig. 5D**). We also found overlapping upregulation of migration-associated genes such as *SEC61G* and *MCRIP1* ^74,75^. The 13 HARs we considered here are not located near each other in the genome and do not share gene targets, supporting that the convergent patterns we observed due to HAR perturbations are the result of independent dysregulation of genes that participate in a common regulatory network.

**Figure 5.**
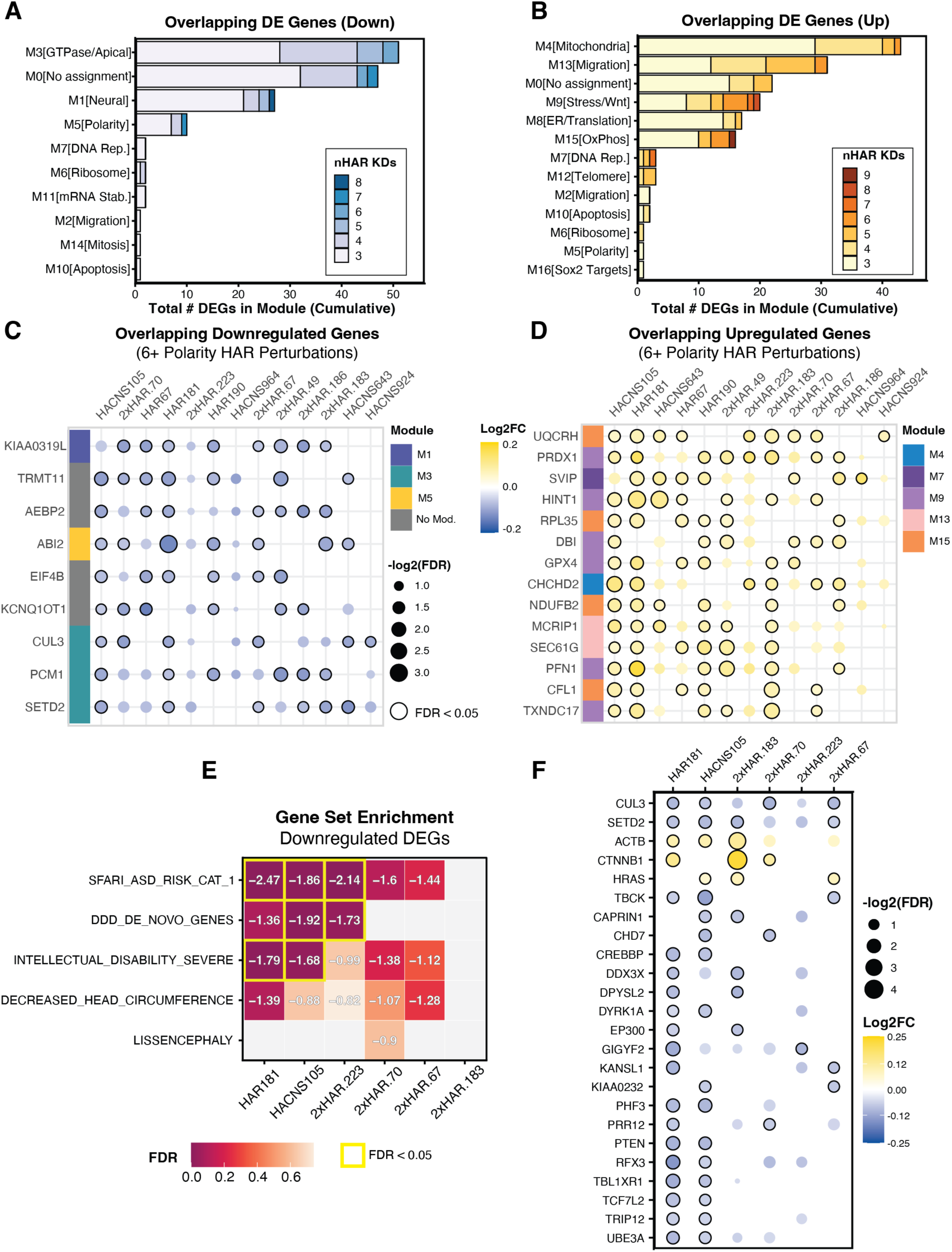
HAR perturbations converge on overlapping sets of differentially expressed genes and are enriched for neurodevelopmental disorder risk genes. **A-B.** Module membership of genes downregulated (A) or upregulated (B) by 3 or more perturbations within the subset of 13 HARs affecting polarity modules. The X-axis is the cumulative total number of DEGs; the color indicates the subsets of DEGs affected by the number of HAR perturbations indicated in the inset legend. **C-D.** Genes downregulated (C) or upregulated (D) by at least 6 separate perturbations affecting polarity-associated modules. Gene names are plotted on the X-axis while HAR perturbations are plotted on the Y-axis. The color of each circle indicates the log2 fold change in expression of each gene and the size of each circle indicates the significance (-log2(FDR)); significant changes are outlined in black. Module membership for each gene is indicated by the ribbon at the left of each plot; color assignments for each module are shown in the legend. **E.** Results from pre-ranked Gene Set Enrichment Analysis for the 6 HAR perturbations downregulating both modules M3 and M5. DEG sets resulting from these perturbations were tested for enrichment of NDD-associated genes. Normalized Enrichment Scores are reported in each box. Significant enrichments (FDR < 0.05) are highlighted in yellow. Grey boxes indicate cases where the NDD gene set had fewer than 15 genes in common with the DEG gene set. **F.** ASD-associated genes affected by 2 or more of the 6 HAR perturbations in (E). The X-axis shows SFARI Category 1 ASD-risk genes and HAR perturbations are plotted on the Y-axis. Changes in expression for each gene are labeled as in **C** and **D**.

### HAR perturbations affecting polarity modules dysregulate genes associated with risk for neurodevelopmental disorders

We noted that DEGs in perturbations affecting apicobasal polarity gene expression included genes associated with neurodevelopmental disorders, such as *SETD2* and *CUL3*. This raised the possibility that HARs may be more broadly involved in the regulation of NDD-associated genes, as has been previously sugested^76^. To assess this, we tested for overrepresentation of NDD-associated genes among DEGs detected for the 6 HAR perturbations that downregulated the expression of both polarity-associated modules (M3 and M5). Using Gene Set Enrichment Analysis (GSEA) on ranked lists of DEGs for these perturbations, we calculated enrichments for SFARI Autism Spectrum Disorder (ASD) Risk Category 1 genes^77^, genes identified as *de novo* mutations in developmental disorders^78^, genes associated with schizophrenia^79^, and genes associated with intellectual disability, decreased head circumference, lissencephaly, or dyslexia (Methods).

We found that downregulated DEGs resulting from 3 of the 6 HAR perturbations (HAR181, *HACNS105*, 2xHAR.223) were enriched for genes in the SFARI ASD Risk Category 1 gene set. Downregulated DEGs resulting from 2 of the 6 HAR perturbations (*HACNS105*, 2xHAR.223) were similarly enriched for genes with *de novo* mutations associated with developmental disorders, and DEGs resulting from 2 perturbations (HAR181, *HACNS105*) were enriched for severe intellectual disability-associated genes (**Fig. 5E**). Genes implicated in lissencephaly were enriched among genes upregulated due to perturbation of *HACNS105* (**Table S11**). We found no enrichment among upregulated or downregulated genes for schizophrenia-associated or dyslexia-associated genes. Twenty-four differentially expressed genes associated with ASD risk were common to 2 or more perturbations, the majority of which were downregulated, including *CUL3*, which maintains cytoskeleton protein homeostasis in NSCs^80^ (**Fig. 5F**). Three ASD risk genes were seen to be upregulated by multiple perturbations, including *CTNNB1*, *ACTB*, and *HRAS*, all of which are associated with cell migration processes^81–83^. Thus, these 6 HARs appear to influence the expression of overlapping sets of genes implicated in NDD risk.

### HAR181 perturbation disrupts rosette formation in cultured NSCs

Our results suggested that some HAR perturbations would lead to disruption of NSC apicobasal polarity, a feature required for proper rosette formation. Rosettes are cell groupings which form spontaneously in monolayer NSC culture as a result of intrinsic polarity signaling^84^. Cells within rosettes exhibit asymmetric localization of PARD3 at the center of the cell grouping, where the NSCs form adherens junctions with one another in a manner resembling the specialized apical surface of aRG^84–86^ (**Fig. 4B**). We therefore hypothesized that silencing HAR181, which had the strongest effects on polarity module expression, may result in disruption of rosette morphology, leading to a disordered cell grouping morphology in cultured NSCs. To test this, we transfected KRAB:dCas9-expressing NSCs with an sgRNA targeting HAR181, its target gene *CADM1*, or a NTC sgRNA. We visualized rosette formation after 7 days of NSC induction with immunofluorescence staining for NES and PARD3.

Representative images of cell groupings, defined as arrangements of 4 or more adjoined cells, from each transfection condition are shown in **Figure 6A**. While cells transfected with NTC sgRNAs formed rosettes, cells transfected with sgRNAs targeting HAR181 or *CADM1* largely formed unstructured cell groupings which did not resemble rosettes. To quantify the extent of rosette malformation, immunofluorescence images of NSCs from each condition were scored by 3 independent, blinded evaluators, and categorized as rosette-like or as unstructured cellular aggregates. Scoring criteria are detailed in **Figure S12**. We found that rosette formation was disrupted in HAR181 and *CADM1* knockdown conditions compared to control cells (**Fig. 6B**), and the proportion of rosette structures to unstructured cell groupings was significantly reduced in sgHAR181 and sg*CADM1* conditions compared to NTC (two-way ANOVA followed by Tukey post-hoc HSD test; **Fig. 6C**). The unprocessed scoring results are tabulated in **Table S13**. These results support the prediction from the Perturb-seq screen that HAR181 is a critical regulator of NSC polarity via *CADM1*.

**Fig 6.**
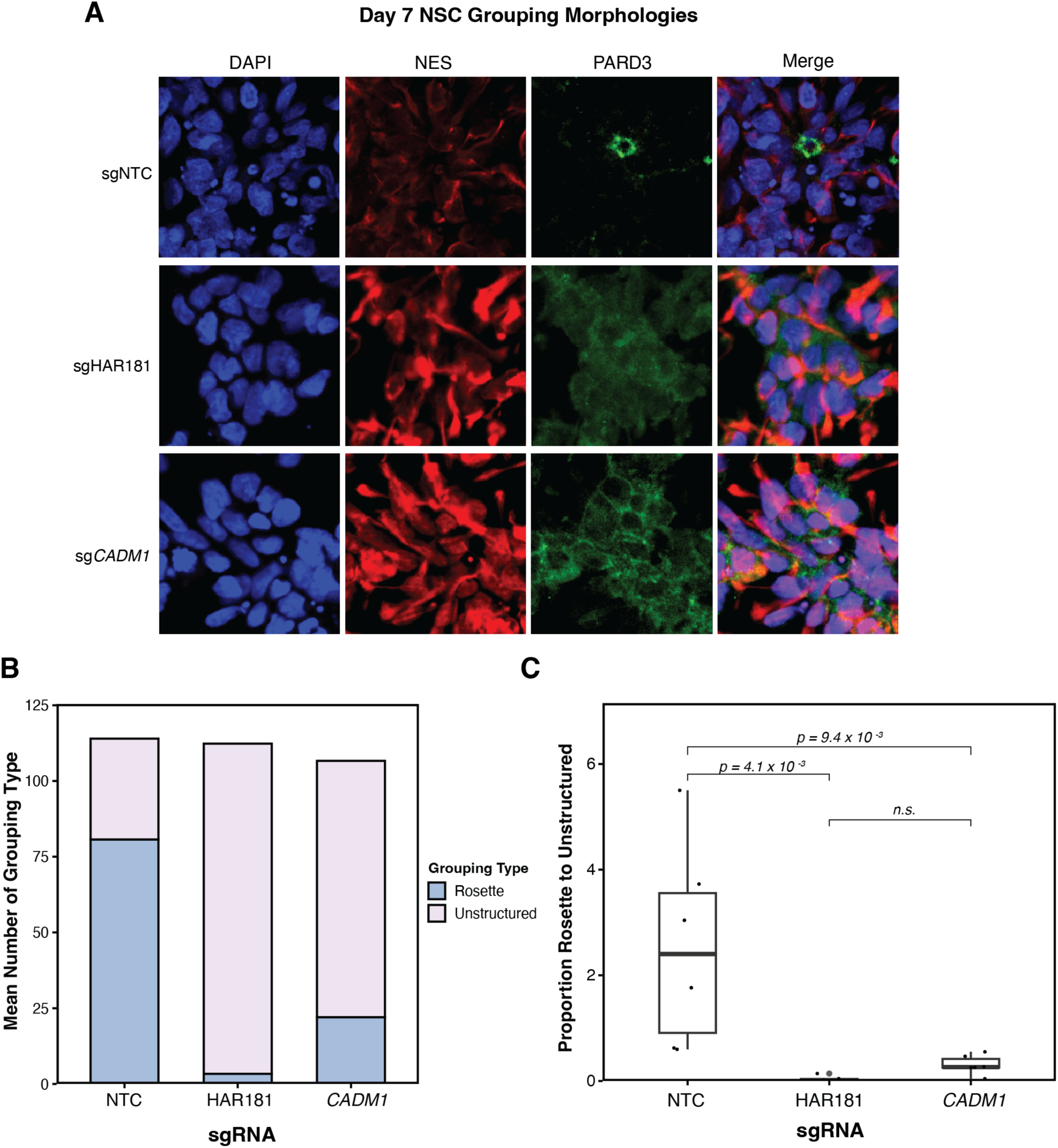
Perturbation of HAR181 disrupts NSC rosette formation. **A.** Representative images of day 7 NSCs transfected with a non-targeting sgRNA (NTC), an sgRNA targeting HAR181 (sgHAR181), and an sgRNA targeting *CADM1* (sg*CADM1*), taken 4 days post-transfection. Localization of PARD3 is shown in green, NES is shown in red and DAPI nuclear staining is shown in blue. **B.** Quantification of the number of rosette-like cell groupings and unstructured cell groupings in each condition. Quantification was performed by 3 independent scorers blinded to the sgRNAs used. Mean counts for each type of cell grouping (rosette-like or unstructured) are shown as stacked bars. **C.** The ratio of rosette-like groupings to unstructured groupings in each condition. Significance was calculated via two-way ANOVA (with the sgRNA and replicate as variables), followed by Tukey post-hoc Honest Significant Difference tests across sgRNA treatments.

## Discussion

Here we used a Perturb-seq screening approach to interrogate the regulatory functions of 180 Human Accelerated Regions in human NSCs. To date, biological characterization of HARs has been limited to a handful of loci. Our data thus advance understanding of the transcriptome-wide regulatory roles of HARs in neurodevelopment and the biological processes in which they operate. Further, though convergent functions of HARs have been hypothesized, no previous study has demonstrated overlapping regulatory roles for HARs in a parallelized assay. We find that multiple HAR perturbations result in the repression of gene modules associated with apicobasal polarity, and demonstrate that the silencing of one HAR, HAR181, disrupts NSC polarity and rosette morphology. The regulation of apicobasal polarity in NSCs is crucial to the generation of bRGs and cortical expansion in development, and is implicated in human cortical evolution^26,49^. As a role for HARs in specifying bRG fate has never been identified, these results provide new insight into the functions of HARs in cortical development.

Our approach relied on direct-capture Perturb-seq to quantify genome-wide transcriptional changes as a result of enhancer perturbation. Enhancer perturbation screens pose additional challenges compared to gene-targeting screens, which employ analytical approaches that presume single, discrete gene targets. Consistent with other studies, we gained valuable insights into the global transcriptional effects of HAR perturbation by focusing on gene co-expression models in addition to individual genes. This approach resulted in testable biological predictions for the functions of multiple HARs in discrete processes. We confirmed one of these predictions: HAR181 repression resulted in the downregulation of polarity-associated genes, mislocalization of PARD3, and the disruption of rosette formation in culture. These results support the validity of our analytical approach. Other HARs with significant effects in our screen appear to have similar biological roles, yielding additional candidates for future experimental studies.

It has previously been postulated that the regulatory functions of different HARs may converge on particular genes and pathways ^5,17,21^. Our design allowed us to assess the global transcriptional effects of multiple HAR perturbations in a single, internally controlled assay, providing us a means to interrogate this hypothesis. Previous predictions of convergent HAR function were based on observations that their known or putative gene targets were enriched for particular neurodevelopmental functions. Here we found convergent dysregulation of gene modules from HARs without shared targets. This indicates that HARs may share another layer of convergence, namely regulation of the same gene networks via the action of independent *cis*-regulatory targets. Among the most significantly downregulated gene networks were GTPase/apical domain- and polarity-associated modules, which were convergently downregulated by 13 HAR perturbations. *In vivo*, the regulation of NSC polarity has direct consequences on cortical expansion in neurodevelopment, as the establishment of the apical domain in aRG limits expansion of the proliferative niche until the emergence and basal migration of the bRG, which lack an apical domain^2,46,49^. These perturbations also exhibited transcriptional effects associated with increased metabolism, migration, and stress. Further, beyond dysregulation of common gene co-expression networks, HAR perturbations also converged on the downregulation of particular genes. Among these was *SETD2*, a chromatin remodeler and cytosolic modifier of tubulin which has previously been shown to contribute to neuronal cell polarity, and whose knockdown has been found to contribute to increased translational output, migratory gene expression, and disrupted NSC lamination, phenotypes which are consistent with our results^66–69,87,88^. Another gene downregulated by several HAR perturbations was *PCM1*, and important regulatory of epithelial cell structure and the timing of neurogenesis, as well as mitotic spindle orientation, a process known to coordinate the emergence of basal cortical progenitors from aRG^71,89^. We therefore posit that these genes may serve as possible nexuses of trans-regulation for multiple HARs, mediating HAR gene targets and their shared downstream effects on polarity networks.

HARs have been shown to encode human-specific regulatory functions during neurodevelopment. One important consideration for our study is that the CRISPR repression approach we used did not directly test the human-specific functions of HARs, as a reduction in activity does not necessarily model the ancestral state of the region. ^15,22^Human-specific substitutions in HARs may contribute to increased or decreased gene expression, alterations in the binding of or response to transcription factors, shifts in the timing of gene activation or inactivation, among other possibilities. We also did not directly compare the effect of perturbing HARs and their chimpanzee orthologs. However, in light of our results, we can now place HARs into specific neurodevelopmental pathways and gene co-expression networks. This provides the basis for biologically informed studies of human-specific functional changes in HARs and the traits that they may have influenced.

Our result that HARs regulate NSC apicobasal polarity has implications in human cortical evolution. The expanded proliferative niche of the bRG progenitor population can ultimately produce more neurons than the more restricted aRG population, and gyrencephalic species, including humans, have been shown to have more basal progenitors in the developing cortex^90–94^. Thus, the evolutionary expansion of the bRG population, either by increased asymmetric divisions of aRGs or by increased symmetric divisions of bRGs, has been one proposed cellular mechanism for human cortical expansion^26,49,95,96^. Our findings suggest that the repression of HARs by CRISPRi leads to a repression of genes associated with the apical domain, that is, genes known to be expressed in aRG. This may reflect an ancestral role for these HARs in the control of apicobasal polarity that is likely shared between human and other primate species. However, human-specific changes in HAR function may alter the level and timing of expression of polarity-associated genes, potentially driving reduced expression at some point during cortical development that would promote delamination and subsequent proliferation of bRGs. Further experimentation and comparative analyses across species will be required to test this putative mechanism.

Our findings establish previously unknown regulatory functions for HARs in gene networks associated with apicobasal polarity, a crucial process in cortical development and evolution. Our data further indicate that HARs act convergently to contribute to the expression of these networks via independent *cis*-regulatory targets. We support our approach by demonstrating that the repression of HAR181 or *CADM1* disrupts NSC polarity and rosette formation in culture. More work will be necessary to discern the role of human-specific changes in HAR function in these processes. Understanding the evolutionary mechanisms of HARs in neurodevelopment requires the means to generate strong hypotheses and identify candidate loci, and our study has provided a basis for these future studies.

## Acknowledgements

We thank Y. Gilad and I. Gallego Romero for generating and providing the human iPSC lines used in this study; Z. Smith for technical assistance and support with single-cell partitioning; S. de la Cruz Molina for experimental advice; B. Lesch, I-H. Park and S. Reilly for manuscript advice. We thank Yale Flow Cytometry for their assistance with FACS; the core supported in part by an NCI Cancer Center Support Grant # NIH P30 CA016359. This work was supported by an award from the Eunice Kennedy Shriver National Institute of Child Health and Human Development (NICHD; R01 HD102030, to J.P.N); NSF Graduate Research Fellowships (to M.A.N., M.M. and K.M.Y.); and an NICHD F32 Postdoctoral Fellowship (F32 HD108935, to M.B). This research program and related results were also made possible by the support of the NOMIS foundation (to J.P.N.).

## Author contributions

M.A.N. and J.P.N conceived and designed the study. M.A.N. optimized cell culture protocols, performed direct-capture Perturb-seq, generated and sequenced libraries, performed CRISPR validation experiments. M.A.N. and Y.J. performed immunofluorescence microscopy. M.A.N., K.M.Y., and M.M. conducted computational analysis. J.W.Y. provided assistance in cell culture. K.M.Y. and M.B. provided advice on statistical approaches and visualizations. J.W.Y., M.M. and R.A-S. scored immunofluorescence images. M.A.N. and J.P.N. wrote the manuscript with input from all authors.

## Data and code availability

All data generated in this study is available at the Gene Expression Omnibus under accession number GSE270828.

All original code generated for this study is available at GitHub: https://github.com/NoonanLab/Noble_et_al_2024 and Zenodo: https://doi.org/10.5281/zenodo.12600310

## Methods

### Experimental Approach

#### Cell lines

Feeder-free human iPSC lines H20961, H23555, and H28126 were provided by Dr. Yoav Gilad. All lines were maintained with mTeSR (STEMCELL Technologies catalog #85850) on 6-well plates coated with Gibco Geltrex hESC-qualified Basement Memebrane (ThermoFisher cat #A1413301) using previously established maintenance protocols^97^. Cells were maintained in a 37 °C incubator with 5% CO_2_. Normal karyotypes for each line were confirmed, and cells were confirmed to be free of mycoplasma contamination.

#### CRISPR sgRNA library design

We aimed to target a subset of the 1592 annotated HARs identified from a set of key studies^9–11^. To target the sgRNA library to neurodevelopmentally active regions, HARs were scored and ranked based on their representation in public datasets. The genomic coordinates of HARs were intersected with activating chromatin marks in neural tissue from the NIH Roadmap Epigenome Mapping Consortium^98^, chromatin accessibility datasets from human brain organoids^33,99,100^, and proliferation phenotype data from CRISPR screening in NSCs^18^. We additionally scored the potential for species-specific activity using comparative ChIP data between human and rhesus macaque brain tissue, as well as comparative chromatin accessibility between human and chimpanzee brain organoids^32,33^.

Our initial aim in designing sgRNAs for HARs was to produce a set of high-quality sgRNAs which could be used in both human and chimpanzee cells with limited background-specific effects. We utilized FlashFry^101^, a command-line tool which generates a database of possible sgRNAs in any genome, while also generating specificity metrics for these potential sgRNAs. Using this, we generated a database of all possible sgRNA binding sites for both the human Hg38 and chimpanzee PanTro6 genomes. Our inputs for sgRNA discovery were the top 600 ranked HARs with 300bp flanking extensions, filtered to be at least 2kb away from gene promoters. We then filtered sgRNAs which had potential off-target binding sites with 2 or fewer mismatches and sgRNAs with an MIT specificity score of below 75^102^. Using machine-learning-based metrics from CRISPRi experiments^103^, we removed out sgRNAs with a predicted on-target score of below 0.40 and on off-target score of above 0.50. We only retained sgRNAs which identical sequences between human and chimpanzee. Further, we filtered out sgRNAs which had differences of MIT specificity scores greater than 5, CRISPRi-based on-target score differences of greater than 0.15, and CRISPRi-based off-target score differences of greater than 0.20. Finally, we selected the two highest scoring sgRNAs per target region whose centers did not overlap within 10bp. For two highly-ranked HARs of interest, *HACNS10* and *HACNS968*, no pair of sgRNAs met these criteria, so their two highest-scoring sgRNAs were selected. Additionally, 5 non-targeting sgRNA control sequences were obtained from a previous study^18^. We also included 1 scrambled sgRNA designed by Sigma Millipore. Gene-targeting positive control sequences were obtained from a previous study^34^. We used one enhancer sequence proximal to *DLGAP1* which elicited strong phenotypes in Geller et al., 2024 as an additional positive control.

The production of lentiviral particles containing sgRNA protospacer sequences compatible with feature barcoding for CRISPR was performed by Sigma Millipore. Lentiviral sgRNA constructs included a direct-capture sequence adjacent to the protospacer, so that sgRNAs are captured and affixed with cell-barcodes alongside mRNA transcripts during single-cell partitioning. The U6-driven sgRNA-expression vector contained the 10x Capture Sequence 1 integrated in the 3’ end of the protospacer and a BFP marker.

#### CRISPRi line generation

The iPSC lines were transduced with KRAB:dCas9 + BSR lentiviral particles (CRISPRIE, was provided by Sigma Millipore with the custom lentiviral pool) at an MOI of 0.10. Each line underwent an antibiotic (Blasticidin S hydrochloride; Thermo catalog #J67216.8) selection kill curve to determine the concentration which killed all wild-type cells within 7 days (between 2.5 and 5 µg/µL). Transduced cells were selected for 7 days with regular passaging, then expanded on antibiotic-free mTeSR, resulting in polycolonal CRISPRi-competent lines. After each freeze-thaw cycle, cells were re-selected with Blasticidin for 72h to ensure transgene expression. We validated expression of *Oct4*, *Sox2*, and *Nanog* in transduced iPSCs to confirm pluripotency marker expression. We measured dCas9 expression by qPCR in both iPSC and NSC states to confirm continued expression of dCas9 through differentiation. Perturbation experiments were performed without the addition of selection antibiotics.

#### Neural stem cell differentiation from iPSCs

To induce the NSC state, iPSCs were released with Accutase (STEMCELL Technologies cat #07920) and plated at a high density of (250kcells/cm^2^ in 6-well plates) as a single-cell suspension in StemDiff Neural Induction Media with Dual SMAD Inhibition (STEMCELL Technologies cat #08581), supplemented with 10 µM ROCK inhibitor Y-27632 (STEMCELL Technologies cat #72304). Plates were coated with Gibco Geltrex Growth Factor Reduced Basement Membrane (Thermo catalog # A1413202). Cells were maintained at high density for 3 days with daily media changes and addition of ROCK inhibitor. Cells were then passaged at a density of 250k cells/cm^2^ when confluent, adding ROCK inhibitor after each passage. Assays were performed after 9 days of dual SMAD inhibition.

#### CRISPRi knockdown efficiency validation

To verify detectable reduction of target gene expression in our polyclonal iPSC-derived NSCs, we differentiated transgenic human line H23555-KRAB with dual SMAD inhibition media as described above. After 3 days of differentiation, cells were transfected via Amaxa Mouse NSC Nucleofection Kits (Lonza catalog #VPG-1004) with a BFP-marked sgRNA expression cassette (Addgene catalog #107722) containing an sgRNA targeting the *GRN* promoter, or containing a non-targeting control sgRNA. After 48 hours, BFP expression was assessed via immunofluorescence imaging. RNA was then extracted for RT-qPCR (see qRT-PCR section). *GRN* expression was normalized to *EEF1* and *TBP* expression, and knockdown efficiency was calculated relative to the rate of transfection.

#### Perturb-seq, single-cell library generation, and sequencing

Human KRAB:dCas9 cells were transduced with pooled lentiviral library (MOI 0.30) after 3 days of dual SMADi neural induction, immediately after passaging. After 48 hours of recovery, the cells were sorted by FACS, gating on BFP expression (sgRNA lentiviral marker), and re-plated in differentiation media with 10 µM ROCK inhibitor. Cells continued dual-SMAD inhibition for an additional 4 days.

Cells were then collected for single-cell RNA-seq. Before single-cell partitioning, cells were released with Accutase and resuspended in Dulbecco’s PBS with 0.04 % BSA on ice. Cells were loaded (9,000 cells per GEM lane) into a 10X Chromium Controller using GEMs and reagents for 3’ v3.1 (Dual Index) single cell RNA-sequencing with Feature Barcoding for CRISPR screening (10X Genomics, catalog #CG000316). Reverse-transcription and library preparation proceeded immediately according to the manufacturer’s user guide. Libraries for gene expression and CRISPR-guide capture are both generated for each GEM lane and indexed individually. Fragment amplification steps were performed using an Eppendorf Mastercycler X50s. Fragment sizes were assayed using an Agilent TapeStation with D5000 reagents (Agilent catalog # 5067-5593). Library concentrations were determined using Invitrogen Qubit dsDNA HS reagents (ThermoFisher catalog #Q32851). Libraries were sequenced using an Illumina NextSeq2000, with P3 200 cycle reagents (Illumina catalog # 20040560), to a depth of ∼40,000 reads/cell for gene expression libraries and 5,000 reads/cell for CRISPR guide-capture libraries.

#### Bulk sgRNA validation assays

Individual sgRNA sequences from the screen were clones into an sgRNA expression vector with GFP (Addgene catalog #107721). sgRNAs targeting the promoters of predicted HAR target genes were also designed and cloned. Vectors were prepared for transfection using E.Z.N.A. Endo-free Plasmid DNA Mini Kits II (Omega Bio-Tek catalog # D6950-01). iPSCs lines were differentiated to NSCs for 4 days, then nucleofected with 1ug vector using Amaxa Mouse NSC Nucleofection kits (Lonza catalog #VPG-1004). Transfection efficiency was assessed by quantifying GFP expression via CytoFLEX flow cytometer (Beckman).

#### Quantitative RT-PCR

RNA was extracted from 1-2 million cells per sample using Qiagen RNeasy Plus Kits (catalog #74134). DNA was removed using gDNA eliminator columns and on-column RNase-free DNase (Qiagen catalog #79254). Reverse transcription and single-strand synthesis was performed using SuperScript III First-Strand Synthesis Supermix for qRT-PCR (ThermoFisher catalog #11752250), and RNA was eliminated using provided RNase (Invitrogen catalog #11752-050). We performed RT-PCR using the LightCycler 480 (Roche) with LightCycler 480 SYBR Green I Master (Roche catalog #04887352001). Primers for qPCR spanned exon junctions and were validated using a melting curve to ensure each primer set yielded a single amplicon. CTs were normalized to *EEF1* or *TBP* expression.

#### Immunofluorescence and cell group morphology scoring

After transfection, cells were expanded in 4-chamber Millicell EZ Slides (Milipore catalog #PEZGS0416) coated with Geltrex. After washing with 1x DPBS, cells were fixed with 4% PFA for 15 minutes. Cells were permeabilized with 0.1% TritonX then stained with rabbit anti-SOX2 (Millipore catalog #AB5603; 1:250), mouse anti-Nestin (Millipore catalog #MAB5326; 1:250), and/or rabbit anti-PARD3 (Novus Biologicals, catalog #NBP1-88861; 1:500) overnight at 4 °C. After washing with TritonX and DPBS, samples were incubated with Alexa Fluor 568 donkey anti-mouse IgG (Molecular Probe catalog #A10037; 1:250) and Alexa Fluor 488 donkey anti-rabbit IgG (Molecular Probe catalog #A21206; 1:250) and mounted with DAPI for imaging. Fluorescence images were captured with a Zeiss LSM 880 confocal microscope.

For cell-grouping quantification, equivalent fields of 1870x1870µm were imaged from two independent replicates per condition. Z-stacks were captured in 2µm steps for each image. Image processing and quantification was performed using FIJI^104^. The maximum intensity value of each axis was projected on the 2D plane using the Z project function of FIJI. Image labels were deidentified and randomized for blind scoring by 3 independent evaluators, using the cell grouping morphology criteria described in **Figure S12**. Briefly, cell groupings were defined as 4 or more adjoined cells with junctions marked by PARD3. Morphologies fell into 4 categories, with types 1 and 2 representing highly-structured or mostly-structured rosette-like formations, and types 3 and 4 represented unstructured or disordered cell aggregates. Scoring was performed using the FIJI Cell Counter function. Significance was determined using a 2-way ANOVA with Tukey post hoc HSD tests for the effects of sgRNA condition and replicate on the ratio of rosettes to unstructured grouping morphologies.

### Quantification and Statistical Analysis

#### Single cell RNA-seq processing

Single-cell sequencing reads from the Perturb-seq experiment were demultiplexed using DRAGEN 4.2 (Illumina). Reads were then aligned to the human genome GRChg38 using CellRanger v7.0.1, and cells across batches and GEM lanes were aggregated using the CellRanger aggr—function without down-sampling. Cell barcodes were labeled with GEM-lane suffixes. The the resulting UMI count matrices were processed using the Seurat R package (v4.3.0)^105^. We removed cells with UMI counts below 1000, then identified and removed potential multi-cell doublets using scDblFinder with a predicted doubleting rate of 4.8% based on the loading concentration, partitioning by GEM lane^106^. Cells with UM1 counts below the second standard deviation of the mean for each batch were removed along with cells whose proportion of mitochondrial reads was 10% or greater. We normalized gene expression using the Seurat function NormalizeData with default parameters. The top 2000 highly variable features were subset (FindVariableFeatures) and scaled (ScaleData).

To assign sgRNAs to cells, we calculated sgRNA UMI thresholds as the 99.6th quantile in the distribution of sgRNA UMI counts in filtered cells. sgRNA thresholds were subtracted from the sgRNA count matrix. Thresholds were calculated within batches. sgRNAs were assigned to cells when resulting UMI counts for a given sgRNA were 2x greater than counts for the next most represented sgRNA in that cell. Cells with multiple sgRNA UMI counts above the UMI threshold in proportions less than 0.50 were considered multi-sgRNA cells and removed from the downstream analyses. Only cells with a single assigned sgRNA were retained.

#### Differential expression analysis

Differential expression was calculated using Wilcoxon Rank Sum using the Seurat function FindMarkers (log2FC = 0.05, min.pct = 0.01). Cells assigned sgRNAs for the same target were compared to all cells bearing NTC sgRNAs. The resulting P-values were adjusted using the false discovery rate within each comparison.

#### Weighted gene co-expression network analysis

To define gene co-expression modules in the data, we implemented the hdWGCNA^43^ package to performed weighted gene correlation network analysis across all Seurat clusters of the CRISPRi single-cell data. Only genes with expression in at least 10% of cells were considered. Modules were named according to the number of member genes, with the largest module as M1. The majority of genes were not assigned a module and were labeled as M0. Module expression is summarized using harmonized module eigengenes (hMEs), corresponding to the Harmony batch-corrected value of the first principal component of each module’s member gene expression. Hub genes are defined the genes with the highest correlation to its module’s expression (kME). Gene ontology enrichments for each module were obtained using the online tools ShinyGO (http://bioinformatics.sdstate.edu/go74/) and gProfiler (biit.cs.ut.ee/gprofiler/gost). Enrichments were identified with gProfiler for all modules except module M16, which showed no significant enrichments except with ShinyGO. In each case, GO enrichments were calculated against the background set of all genes expressed in at least 10% of cells.

#### Multiple Linear Regression

We fit a multiple linear regression model for each module, with hME values as the response and nFeatures (nGenes detected in each cell), perturbation (sgRNA target) and batch as variables. Our model measuring ME values as a response to perturbation is of a similar design to one previously described^107^. For our calculations, we used the lm() function in R. The MLR model used the formula:

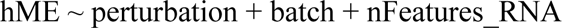

For the perturbation variable, NTC cells were used as the reference, so effect sizes, which are the beta coefficients for each term, can be understood as the estimated shift in the mean ME value given a perturbation. P-values for each perturbation were obtained for each module, and the false discovery rate was calculated from the set of P-values for each module’s MLR model.

#### Gene Set Enrichment Analysis

To perform GSEA, we used the GSEA 4.3.2 utility via MSigDB^108^. Differentially expressed genes for the 6 HARs significantly impacting M3 and M5 were ranked based on the calculation:

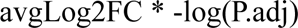

We tested gene sets for severe intellectual disability, primary microcephaly, dyslexia, lissencephaly, and decreased head circumference obtained from MSigDB human phenotype datasets (https://www.gsea-msigdb.org/gsea/msigdb/human/collections.jsp). Gene sets for ASD risk were composed of SFARI category 1 risk genes^77^. Gene sets for schizophrenia risk were obtained of associated rare coding variants^79^. Gene sets for developmental disabilities were the highest significance de novo mutations described from the Deciphering Developmental Disorders consortium^78^. Nominal p-values for enrichment scores were calculated against a null p-distribution obtained after 1000 random permutations of phenotype labels. Enrichment scores were then normalized by gene set size and p-values were adjusted using the false discovery rate.

## References

1. Götz, M. & Huttner, W. B. The cell biology of neurogenesis. Nat. Rev. Mol. Cell Biol. 6, 777–788 (2005).

2. Merino, F. & Götz, M. Neocortical Neurogenesis in Development and Evolution. 1–18 (2023) doi:10.1002/9781119860914.ch1.

3. Mora-Bermúdez, F. & Huttner, W. B. What Are the Human-Specific Aspects of Neocortex Development? Front. Neurosci. 16, 878950 (2022).

4. Levchenko, A., Kanapin, A., Samsonova, A. & Gainetdinov, R. Human accelerated regions and other human-specific sequence variations in the context of evolution and their relevance for brain development. Genome Biol. Evol. 10, evx240- (2017).

5. Won, H., Huang, J., Opland, C. K., Hartl, C. L. & Geschwind, D. H. Human evolved regulatory elements modulate genes involved in cortical expansion and neurodevelopmental disease susceptibility. Nat. Commun. 10, 2396 (2019).

6. Pizzollo, J., Zintel, T. M. & Babbitt, C. C. Differentially Active and Conserved Neural Enhancers Define Two Forms of Adaptive Noncoding Evolution in Humans. Genome Biol. Evol. 14, evac108 (2022).

7. Fair, T. & Pollen, A. A. Genetic architecture of human brain evolution. Curr. Opin. Neurobiol. 80, 102710 (2023).

8. Pollard, K. S. et al. An RNA gene expressed during cortical development evolved rapidly in humans. Nature 443, 167–172 (2006).

9. Pollard, K. S. et al. Forces Shaping the Fastest Evolving Regions in the Human Genome. PLoS Genet. 2, e168 (2006).

10. Prabhakar, S., Noonan, J. P., Pääbo, S. & Rubin, E. M. Accelerated Evolution of Conserved Noncoding Sequences in Humans. Science 314, 786–786 (2006).

11. Lindblad-Toh, K. et al. A high-resolution map of human evolutionary constraint using 29 mammals. Nature 478, 476–482 (2011).

12. Prabhakar, S. et al. Human-Specific Gain of Function in a Developmental Enhancer. Science 321, 1346–1350 (2008).

13. Capra, J. A., Erwin, G. D., McKinsey, G., Rubenstein, J. L. R. & Pollard, K. S. Many human accelerated regions are developmental enhancers. Philos. Trans. R. Soc. B: Biol. Sci. 368, 20130025 (2013).

14. Boyd, J. L. et al. Human-Chimpanzee Differences in a FZD8 Enhancer Alter Cell-Cycle Dynamics in the Developing Neocortex. Curr. Biol. 25, 772–779 (2015).

15. Uebbing, S., et al. Massively parallel discovery of human-specific substitutions that alter enhancer activity. Proc. Natl. Acad. Sci. 118, e2007049118 (2021).

16. Girskis, K. M. et al. Rewiring of human neurodevelopmental gene regulatory programs by human accelerated regions. Neuron 109, 3239–3251.e7 (2021).

17. Whalen, S. & Pollard, K. S. Enhancer Function and Evolutionary Roles of Human Accelerated Regions. Annu. Rev. Genet. 56, 423–439 (2022).

18. Geller, E. et al. Massively parallel disruption of enhancers active in human neural stem cells. Cell Rep. 43, 113693 (2024).

19. Won, H. et al. Chromosome conformation elucidates regulatory relationships in developing human brain. 538, (2016).

20. Keough, K. C. et al. Three-dimensional genome rewiring in loci with human accelerated regions. Science 380, eabm1696 (2023).

21. Pal, A. et al. Resolving the three-dimensional interactome of Human Accelerated Regions during human and chimpanzee neurodevelopment. (2024) doi:10.1101/2024.06.25.600691.

22. Whalen, S. et al. Machine learning dissection of human accelerated regions in primate neurodevelopment. Neuron 111, 857–873.e8 (2023).

23. Replogle, J. M. et al. Combinatorial single-cell CRISPR screens by direct guide RNA capture and targeted sequencing. Nat. Biotechnol. 38, 954–961 (2020).

24. Morris, J. A. et al. Discovery of target genes and pathways at GWAS loci by pooled single-cell CRISPR screens. Science 380, eadh7699 (2023).

25. Alda-Catalinas, C. et al. Mapping the functional impact of non-coding regulatory elements in primary T cells through single-cell CRISPR screens. Genome Biol. 25, 42 (2024).

26. Arai, Y. & Taverna, E. Neural Progenitor Cell Polarity and Cortical Development. Front. Cell. Neurosci. 11, 384 (2017).

27. Replogle, J. M. et al. Combinatorial single-cell CRISPR screens by direct guide RNA capture and targeted sequencing. Nat. Biotechnol. 38, 954–961 (2020).

28. Replogle, J. M. et al. Mapping information-rich genotype-phenotype landscapes with genome-scale Perturb-seq. Cell 185, 2559–2575.e28 (2022).

29. Sunshine, S. et al. Systematic functional interrogation of SARS-CoV-2 host factors using Perturb-seq. Nat. Commun. 14, 6245 (2023).

30. Liu, S. J. et al. Identifying Gene-Treatment Interactions and Targetable Radiation Vulnerabilities in Glioblastoma through Coupling of In Vivo CRISPR Perturbation and Single Cell Transcriptomics. Int. J. Radiat. Oncol.Biol.Phys. 117, S102 (2023).

31. Doench, J. G. Am I ready for CRISPR? A user’s guide to genetic screens. Nat. Rev. Genet. 19, 67–80 (2018).

32. Reilly, S. K. et al. Evolutionary changes in promoter and enhancer activity during human corticogenesis. Science 347, 1155–1159 (2015).

33. Kanton, S. et al. Organoid single-cell genomic atlas uncovers human-specific features of brain development. Nature 574, 418–422 (2019).

34. Tian, R. et al. CRISPR Interference-Based Platform for Multimodal Genetic Screens in Human iPSC-Derived Neurons. Neuron 104, 239–255.e12 (2019).

35. Götz, M., Stoykova, A. & Gruss, P. Pax6 Controls Radial Glia Differentiation in the Cerebral Cortex. Neuron 21, 1031–1044 (1998).

36. Ellis, P. et al. SOX2, a Persistent Marker for Multipotential Neural Stem Cells Derived from Embryonic Stem Cells, the Embryo or the Adult. Dev. Neurosci. 26, 148–165 (2004).

37. Shimizu, F., Watanabe, T. K., Shinomiya, H., Nakamura, Y. & Fujiwara, T. Isolation and expression of a cDNA for human brain fatty acid-binding protein (B-FABP)1The nucleotide sequence reported in this paper have been deposited in the DDBJ, EMBL and GeneBank databases under accession number D88648.1. Biochim. Biophys. Acta (BBA) - Gene Struct. Expr. 1354, 24–28 (1997).

38. Eze, U. C., Bhaduri, A., Haeussler, M., Nowakowski, T. J. & Kriegstein, A. R. Single-cell atlas of early human brain development highlights heterogeneity of human neuroepithelial cells and early radial glia. Nat. Neurosci. 24, 584–594 (2021).

39. Papalexi, E. et al. Characterizing the molecular regulation of inhibitory immune checkpoints with multimodal single-cell screens. Nat. Genet. 53, 322–331 (2021).

40. Bock, C. et al. High-content CRISPR screening. Nat. Rev. Methods Prim. 2, 8 (2022).

41. Jin, X. et al. In vivo Perturb-Seq reveals neuronal and glial abnormalities associated with autism risk genes. Science 370, (2020).

42. Sun, X., Wang, Z., Chen, X. & Shen, K. CRISPR-cas9 Screening Identified Lethal Genes Enriched in Cell Cycle Pathway and of Prognosis Significance in Breast Cancer. Front. Cell Dev. Biol. 9, 646774 (2021).

43. Morabito, S., Reese, F., Rahimzadeh, N., Miyoshi, E. & Swarup, V. hdWGCNA identifies co-expression networks in high-dimensional transcriptomics data. *Cell Rep*. Methods 3, 100498 (2023).

44. Langfelder, P. & Horvath, S. WGCNA: an R package for weighted correlation network analysis. BMC Bioinform. 9, 559 (2008).

45. Werling, D. M. et al. Whole-Genome and RNA Sequencing Reveal Variation and Transcriptomic Coordination in the Developing Human Prefrontal Cortex. Cell Rep. 31, 107489 (2020).

46. Smart, I. H. M., Dehay, C., Giroud, P., Berland, M. & Kennedy, H. Unique Morphological Features of the Proliferative Zones and Postmitotic Compartments of the Neural Epithelium Giving Rise to Striate and Extrastriate Cortex in the Monkey. Cereb. Cortex 12, 37–53 (2002).

47. Yamamoto, H. et al. Impairment of radial glial scaffold-dependent neuronal migration and formation of double cortex by genetic ablation of afadin. Brain Res. 1620, 139–152 (2015).

48. Rakotomamonjy, J. et al. Afadin controls cell polarization and mitotic spindle orientation in developing cortical radial glia. Neural Dev. 12, 7 (2017).

49. Andrews, M. G., Subramanian, L., Salma, J. & Kriegstein, A. R. How mechanisms of stem cell polarity shape the human cerebral cortex. Nat. Rev. Neurosci. 23, 711–724 (2022).

50. Pollen, A. A. & Kriegstein, A. R. Neocortical Neurogenesis in Development and Evolution. 19–39 (2023) doi:10.1002/9781119860914.ch2.

51. Kawaguchi, A. Neuronal Delamination and Outer Radial Glia Generation in Neocortical Development. Front. Cell Dev. Biol. 8, 623573 (2021).

52. Lancaster, M. A. & Knoblich, J. A. Spindle orientation in mammalian cerebral cortical development. Curr. Opin. Neurobiol. 22, 737–746 (2012).

53. Abdi, K. & Kuo, C. T. Laminating the mammalian cortex during development: cell polarity protein function and Hippo signaling. Genes Dev. 32, 740–741 (2018).

54. Nishimura, T. & Takeichi, M. Shroom3-mediated recruitment of Rho kinases to the apical cell junctions regulates epithelial and neuroepithelial planar remodeling. Development 135, 1493–1502 (2008).

55. Hong, E., Jayachandran, P. & Brewster, R. The polarity protein Pard3 is required for centrosome positioning during neurulation. Dev. Biol. 341, 335–345 (2010).

56. Kono, K., Tamashiro, D. A. A. & Alarcon, V. B. Inhibition of RHO–ROCK signaling enhances ICM and suppresses TE characteristics through activation of Hippo signaling in the mouse blastocyst. Dev. Biol. 394, 142–155 (2014).

57. Chou, F.-S., Li, R. & Wang, P.-S. Molecular components and polarity of radial glial cells during cerebral cortex development. Cell. Mol. Life Sci. 75, 1027–1041 (2018).

58. Tavano, S. et al. Insm1 Induces Neural Progenitor Delamination in Developing Neocortex via Downregulation of the Adherens Junction Belt-Specific Protein Plekha7. Neuron 97, 1299–1314.e8 (2018).

59. Penisson, M., Ladewig, J., Belvindrah, R. & Francis, F. Genes and Mechanisms Involved in the Generation and Amplification of Basal Radial Glial Cells. Front. Cell. Neurosci. 13, 381 (2019).

60. Ferent, J., Zaidi, D. & Francis, F. Extracellular Control of Radial Glia Proliferation and Scaffolding During Cortical Development and Pathology. Front. Cell Dev. Biol. 8, 578341 (2020).

61. Grove, M. et al. Abi2-Deficient Mice Exhibit Defective Cell Migration, Aberrant Dendritic Spine Morphogenesis, and Deficits in Learning and Memory. Mol. Cell. Biol. 24, 10905–10922 (2004).

62. Zhang, W. et al. MiRNA-128 regulates the proliferation and neurogenesis of neural precursors by targeting PCM1 in the developing cortex. eLife 5, e11324 (2016).

63. Amano, M., Nakayama, M. & Kaibuchi, K. Rho-kinase/ROCK: A key regulator of the cytoskeleton and cell polarity. Cytoskeleton 67, 545–554 (2010).

64. Govek, E., Hatten, M. E. & Aelst, L. V. The role of Rho GTPase proteins in CNS neuronal migration. Dev. Neurobiol. 71, 528–553 (2011).

65. Ostrem, B. E. L., Lui, J. H., Gertz, C. C. & Kriegstein, A. R. Control of Outer Radial Glial Stem Cell Mitosis in the Human Brain. Cell Rep. 8, 656–664 (2014).

66. Park, I. Y. et al. Dual Chromatin and Cytoskeletal Remodeling by SETD2. Cell 166, 950–962 (2016).

67. Xu, L. et al. Abnormal neocortex arealization and Sotos-like syndrome–associated behavior in Setd2 mutant mice. Sci. Adv. 7, eaba1180 (2021).

68. Kearns, S. et al. Molecular determinants for α-tubulin methylation by SETD2. J. Biol. Chem. 297, 100898 (2021).

69. Xie, X. et al. α-TubK40me3 is required for neuronal polarization and migration by promoting microtubule formation. Nat. Commun. 12, 4113 (2021).

70. Mitchell, B., Thor, S. & Piper, M. Cellular and molecular functions of SETD2 in the central nervous system. J. Cell Sci. 136, (2023).

71. Ge, X., Frank, C. L., Anda, F. C. de & Tsai, L.-H. Hook3 Interacts with PCM1 to Regulate Pericentriolar Material Assembly and the Timing of Neurogenesis. Neuron 65, 191–203 (2010).

72. Neumann, C. A., Cao, J. & Manevich, Y. Peroxiredoxin 1 and its role in cell signaling. Cell Cycle 8, 4072–4078 (2009).

73. Ursini, F. & Maiorino, M. Lipid peroxidation and ferroptosis: The role of GSH and GPx4. Free Radic. Biol. Med. 152, 175–185 (2020).

74. Jin, L., Chen, D., Hirachan, S., Bhandari, A. & Huang, Q. SEC61G regulates breast cancer cell proliferation and metastasis by affecting the Epithelial-Mesenchymal Transition. J. Cancer 13, 831–846 (2022).

75. Ichikawa, K. et al. MCRIP1, an ERK Substrate, Mediates ERK-Induced Gene Silencing during Epithelial-Mesenchymal Transition by Regulating the Co-Repressor CtBP. Mol. Cell 58, 35–46 (2015).

76. Doan, R. N. et al. Mutations in Human Accelerated Regions Disrupt Cognition and Social Behavior. Cell 167, 341–354.e12 (2016).

77. Satterstrom, F. K. et al. Large-Scale Exome Sequencing Study Implicates Both Developmental and Functional Changes in the Neurobiology of Autism. Cell 180, 568–584.e23 (2020).

78. McRae, J. F. et al. Prevalence and architecture of de novo mutations in developmental disorders. Nature 542, 433–438 (2017).

79. Singh, T. et al. Rare coding variants in ten genes confer substantial risk for schizophrenia. Nature 604, 509–516 (2022).

80. Morandell, J. et al. Cul3 regulates cytoskeleton protein homeostasis and cell migration during a critical window of brain development. Nat. Commun. 12, 3058 (2021).

81. Drosten, M. et al. Genetic analysis of Ras signalling pathways in cell proliferation, migration and survival. EMBO J. 29, 1091–1104 (2010).

82. Bunnell, T. M., Burbach, B. J., Shimizu, Y. & Ervasti, J. M. β-Actin specifically controls cell growth, migration, and the G-actin pool. Mol. Biol. Cell 22, 4047–4058 (2011).

83. Wal, T. van der & Amerongen, R. van. Walking the tight wire between cell adhesion and WNT signalling: a balancing act for β-catenin. Open Biol. 10, 200267 (2020).

84. Banda, E. et al. Cell Polarity and Neurogenesis in Embryonic Stem Cell-Derived Neural Rosettes. Stem Cells Dev. 24, 1022–1033 (2015).

85. Grabiec, M. et al. Stage-specific roles of FGF2 signaling in human neural development. Stem Cell Res. 17, 330–341 (2016).

86. Hříbková, H., Grabiec, M., Klemová, D., Slaninová, I. & Sun, Y.-M. Calcium signaling mediates five types of cell morphological changes to form neural rosettes. J. Cell Sci. 131, jcs206896 (2018).

87. Wang, T. et al. SETD2 loss in renal epithelial cells drives epithelial-to-mesenchymal transition in a TGF-β-independent manner. Mol. Oncol. 18, 44–61 (2024).

88. Molenaar, T. M. et al. The histone methyltransferase SETD2 negatively regulates cell size. J. Cell Sci. 135, jcs259856 (2022).

89. Dammermann, A. & Merdes, A. Assembly of centrosomal proteins and microtubule organization depends on PCM-1. J. Cell Biol. 159, 255–266 (2002).

90. Dehay, C., Kennedy, H. & Kosik, K. S. The Outer Subventricular Zone and Primate-Specific Cortical Complexification. Neuron 85, 683–694 (2015).

91. Arai, Y. et al. Neural stem and progenitor cells shorten S-phase on commitment to neuron production. Nat. Commun. 2, 154 (2011).

92. Xing, L., Pinson, A., Mora-Bermúdez, F. & Huttner, and W. B. Neocortical Neurogenesis in Development and Evolution. 137–156 (2023) doi:10.1002/9781119860914.ch8.

93. Kelava, I. et al. Abundant Occurrence of Basal Radial Glia in the Subventricular Zone of Embryonic Neocortex of a Lissencephalic Primate, the Common Marmoset Callithrix jacchus. Cereb. Cortex 22, 469–481 (2012).

94. Benito-Kwiecinski, S. et al. An early cell shape transition drives evolutionary expansion of the human forebrain. Biorxiv 2020.07.04.188078 (2020) doi:10.1101/2020.07.04.188078.

95. Florio, M. & Huttner, W. B. Neural progenitors, neurogenesis and the evolution of the neocortex. Development 141, 2182–2194 (2014).

96. Romero, C. de J., Bruder, C., Tomasello, U., Sanz-Anquela, J. M. & Borrell, V. Discrete domains of gene expression in germinal layers distinguish the development of gyrencephaly. EMBO J. 34, 1859–1874 (2015).

97. Romero, I. G. et al. A panel of induced pluripotent stem cells from chimpanzees: a resource for comparative functional genomics. eLife 4, e07103 (2015).

98. Bernstein, B. E. et al. The NIH Roadmap Epigenomics Mapping Consortium. Nat. Biotechnol. 28, 1045–1048 (2010).

99. Xiang, Y. et al. Fusion of Regionally Specified hPSC-Derived Organoids Models Human Brain Development and Interneuron Migration. Cell Stem Cell 21, 383–398.e7 (2017).

100. Amiri, A. et al. Transcriptome and epigenome landscape of human cortical development modeled in organoids. Science 362, (2018).

101. McKenna, A. & Shendure, J. FlashFry: a fast and flexible tool for large-scale CRISPR target design. Bmc Biol 16, 74 (2018).

102. Hsu, P. D. et al. DNA targeting specificity of RNA-guided Cas9 nucleases. Nat. Biotechnol. 31, 827–832 (2013).

103. Jost, M. et al. Titrating gene expression using libraries of systematically attenuated CRISPR guide RNAs. Nat. Biotechnol. 38, 355–364 (2020).

104. Schindelin, J., et al. Fiji: an open-source platform for biological-image analysis. Nat. Methods 9, 676–682 (2012).

105. Satija, R., Farrell, J. A., Gennert, D., Schier, A. F. & Regev, A. Spatial reconstruction of single-cell gene expression data. Nat. Biotechnol. 33, 495–502 (2015).

106. Germain, P.-L., Lun, A., Meixide, C. G., Macnair, W. & Robinson, M. D. Doublet identification in single-cell sequencing data using scDblFinder. F1000Research 10, 979 (2021).

107. Jin, X. et al. In vivo Perturb-Seq reveals neuronal and glial abnormalities associated with Autism risk genes. Biorxiv 791525 (2019) doi:10.1101/791525.

108. Subramanian, A. et al. Gene set enrichment analysis: A knowledge-based approach for interpreting genome-wide expression profiles. Proc. Natl. Acad. Sci. 102, 15545–15550 (2005).

