## Supplementary_Figures for "Human Accelerated Regions regulate gene networks implicated in apical-to-basal neural progenitor fate transitions"

**Supplementary Materials**

Supplementary Figures


Figure S1: Dual-species design of sgRNA library targeting human accelerated regions
Figure S2: iPSC-derived NSC cell line validation and CRISPRi efficiency
Figure S3: Perturb-seq data processing and quality control
Figure S4: Single-cell CRISPR sgRNA assignment pipeline
Figure S5: Differential expression of HAR perturbations and their predicted targets.
Figure S6: Differential expression and power to detect direct gene perturbations in gene-targeting controls
Figure S7: hdWGCNA module composition, expression correlations, and hub genes
Figure S8: Multiple linear regression of HAR perturbation effects on modules
Figure S9: RT-qPCR of predicted HAR181 and CADM1 targets in independent cell line.
Figure S10: Distribution of overlapping significant DE genes from 1 or more perturbations
Figure S11: Convergence in DE genes among ≥ 4 perturbations affecting polarity modules
Figure S12: Scoring rubric and quantification of NSC cell grouping structures


Supplementary Tables


Table S1: sgRNA library sequences and genomic positions
Table S2: Perturb-seq single-cell metadata
Table S3: HAR perturbation differential expression
Table S4: WGCNA module genes and kMEs
Table S5: Module GO enrichments
Table S6: Single-cell module eigengene values
Table S7: HAR perturbation multiple linear regression results
Table S8: Summary of perturbation effects by module
Table S9: Frequency of differential expression by gene (all perturbations)
Table S10: GSEA NDD gene lists
Table S11: GSEA NDD enrichment results
Table S12: Perturb-seq sgRNA target representation
Table S13: NSC grouping morphology blinded counts

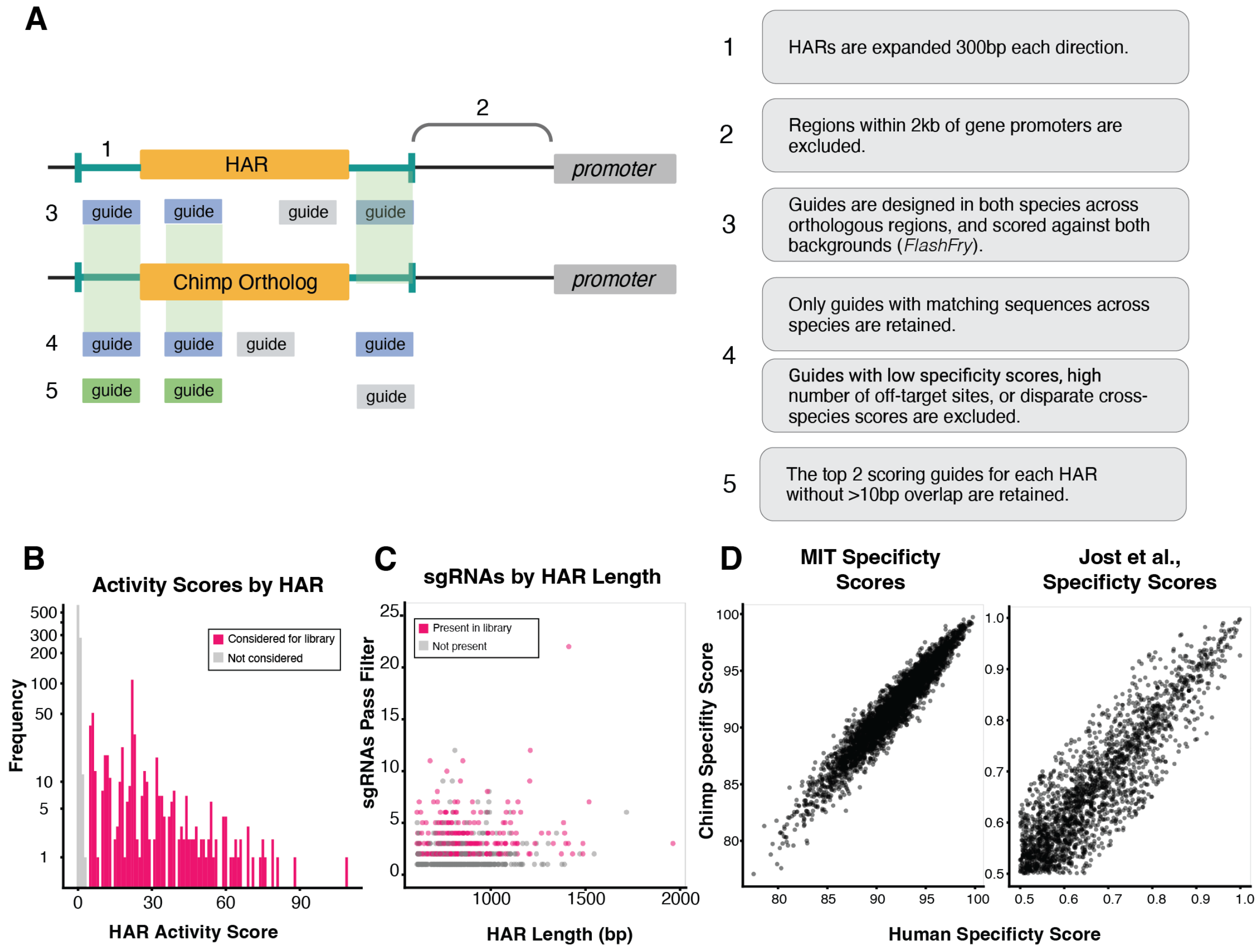


**Fig S1. Dual-species design of sgRNA library targeting human accelerated regions. A.** Schematic detailing sgRNA library design parameters and filtering criteria. The pipeline to generate a high-specificity set of sgRNAs targeting neurodevelopmentally active HARs was optimized allow for its utilization in both human and chimpanzee genomic backgrounds. Guides were stringently filtered using metrics compiled by FlashFry for each background, were required to have identical sequences in human and chimpanzee genomes, and specificity metrics were required to be similar for each background. This figure was made in part using BioRender.com **B.** Histogram of HAR activity scores for 1592 HARs. Each HAR was scored with respect to their intersections with published data for neurodevelopmental activity. We utilized ChIP-Seq data from RoadMap fetal brain tissue and published ChIP-seq datasets, ATAC-seq in organoid studies, proliferation effects from CRISPR screens, and evidence from organoids and tissues for species-biased activity. We then subset the top fraction of HARs with evidence for activity from at least two sources to consider for sgRNA design. **C.** Plot showing the relationship between HAR length and the number of potential sgRNAs which pass QC filters. All possible sgRNAs were initially designed for each active HAR. The number of possible sgRNAs passing QC filters weakly correlated with HAR length. The top two highest scoring, non-overlapping guides (center > 10bp apart) were selected for each HAR, then the top 180 highest scoring HARs were selected to create the final library. Points for sgRNAs which were retained for the final library are highlighted in pink. **D-E.**) Correlations of MIT specificity scores (Hsu et al. 2013) and dCas9 binding specificity scores (Jost et al. 2020) for each possible guide after filtering, comparing scores calculated in both human and chimpanzee genomic backgrounds.

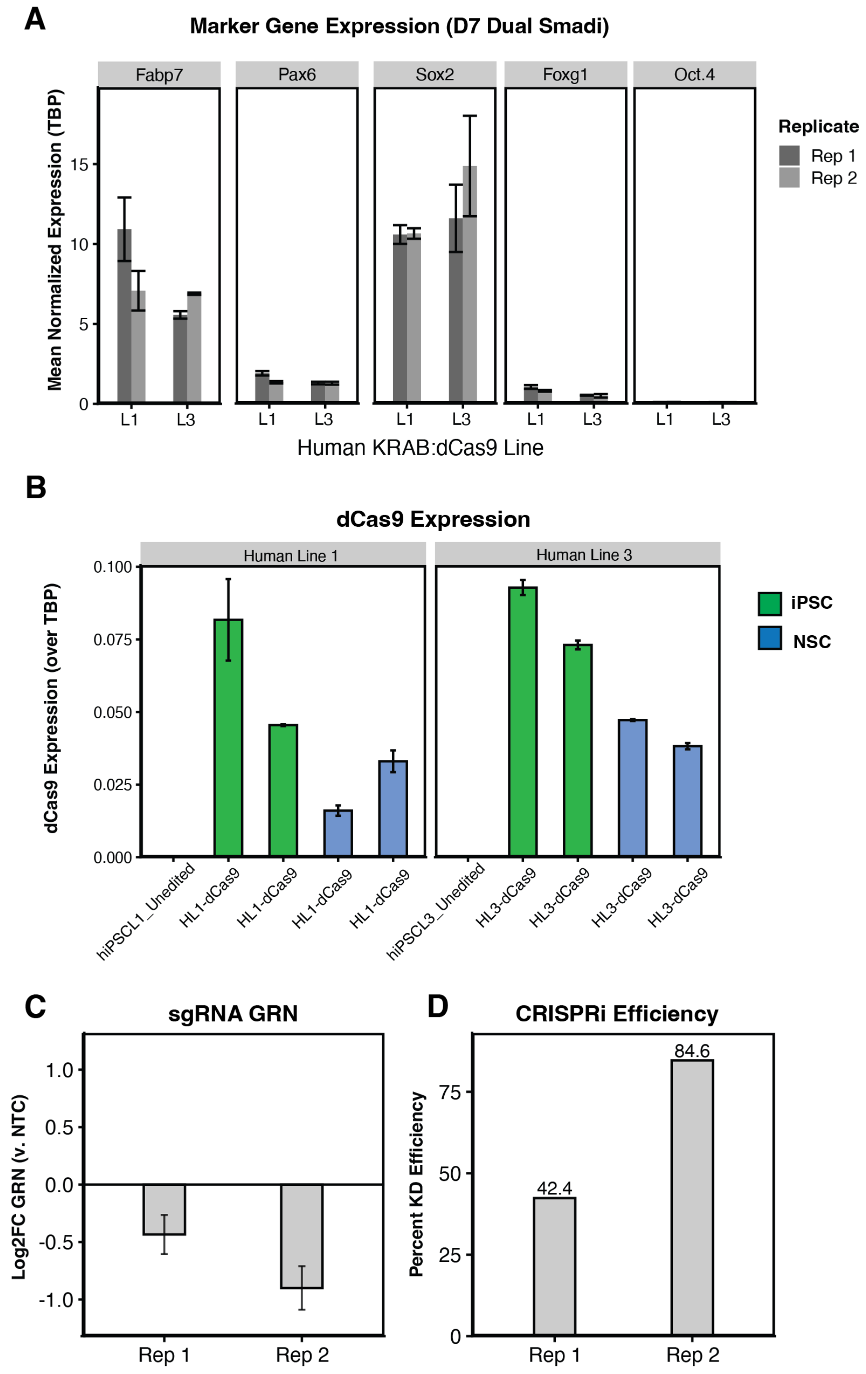


**Fig. S2. iPSC-derived NSC cell line validation and CRISPRi efficiency. A.** RT-qPCR of NSC and iPSC markers from 2 independent human lines (L1 and L2) on day 7 of dual SMAD inhibition. Expression is normalized to *TBP* expression. **B**. RT-qPCR for dCas9 expression in 2 edited human lines while in the iPSC state (green) and NSC state (blue) after 9 days of dual SMAD inhibition. Two replicates are shown. **C**-**D.** Log2 fold change of *GRN* expression from cells transfected with *GRN* targeting sgRNAs compared to cells transfected with NTC sgRNAs. Percent knockdown efficiency was calculated from a theoretical knockdown efficiency equivalent to the transfection rate (number of BFP+ cells over total cells).

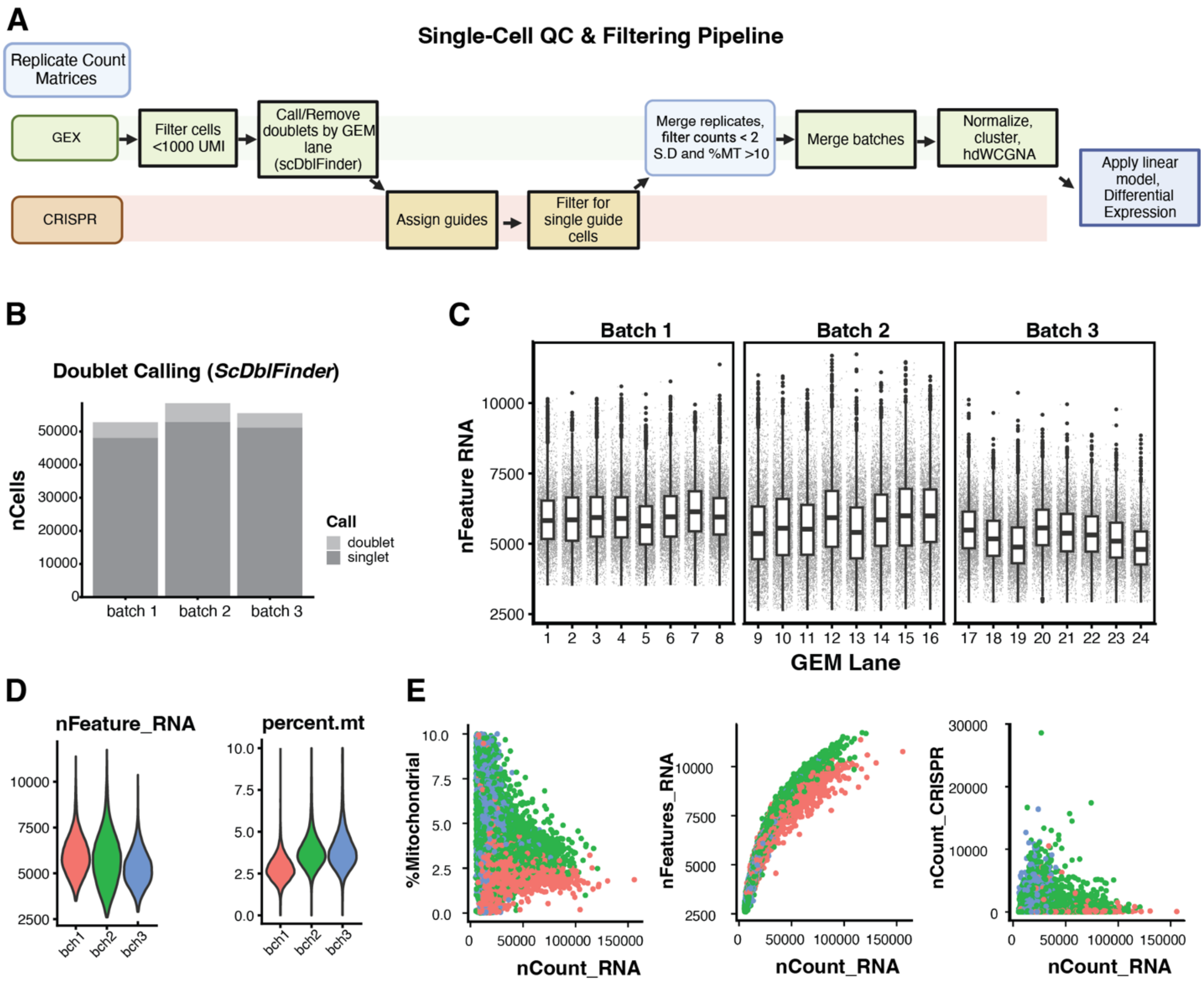


**Fig. S3. Perturb-seq data processing and quality control. A.** Schematic of Perturb-seq filtering and quality control preprocessing steps. Doublets were removed by GEM-lane using scDblFinder and guides were assigned using distributions within batches. Batches are then computationally merged and filtered for quality covariates. This figure was made using BioRender.com. **B.** The proportions of cells removed from each batch after doublet calling using ScDblFinder. **C.** nFeature RNA (number of detected genes) across each GEM lane for each batch. **D**. Violin plot of cell quality covariate distributions, nFeauture_RNA and percent mitochondrial reads, across batches after QC filtering. **E**. Correspondence of total RNA counts (nCount_RNA) to percent mitochondrial reads, nFeatures and CRISPR guide capture counts across batches. Batches are colored as in **D**.

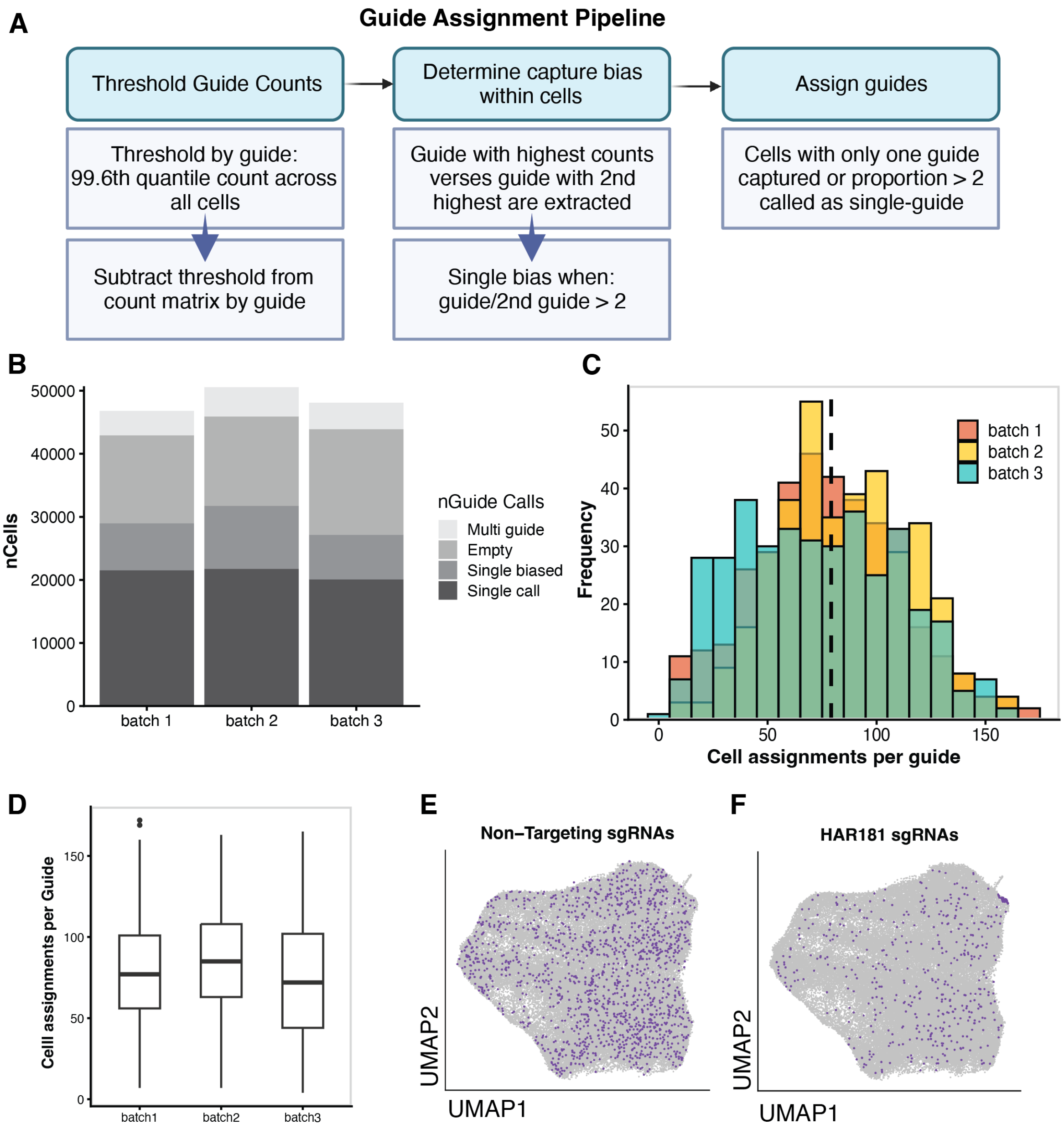


**Fig. S4. Single-cell CRISPR sgRNA assignment pipeline. A.** Schematic of the guide assignment pipeline applied to each Perturb-seq batch. The pipeline accounts for the distribution of each guide across cells, as well as the proportion of counts within each cell. Only cells with a single assigned guide were retained for analysis. This figure was generated using BioRender.com. **B**. The proportions of cells called as single true (only 1 guide detected) or single biased (proportion of 1 guide 2x greater than the next most-detected guide), versus multi-guide calls and cells with no guide passing the filtering step across batches (Methods). **C**. Distribution of guide assignments in cells across batches. **D.** Correspondence of guide-per-cell coverage across batches. **E-F.** UMAPs of gene expression in filtered Perturb-seq cells across all batches, in which cells with NTC sgRNA assignments or HAR181 sgRNA assignments are highlighted. NTC assignments are distributed throughout the data, whereas some HAR181 guide assignments form a small subcluster at the top right.

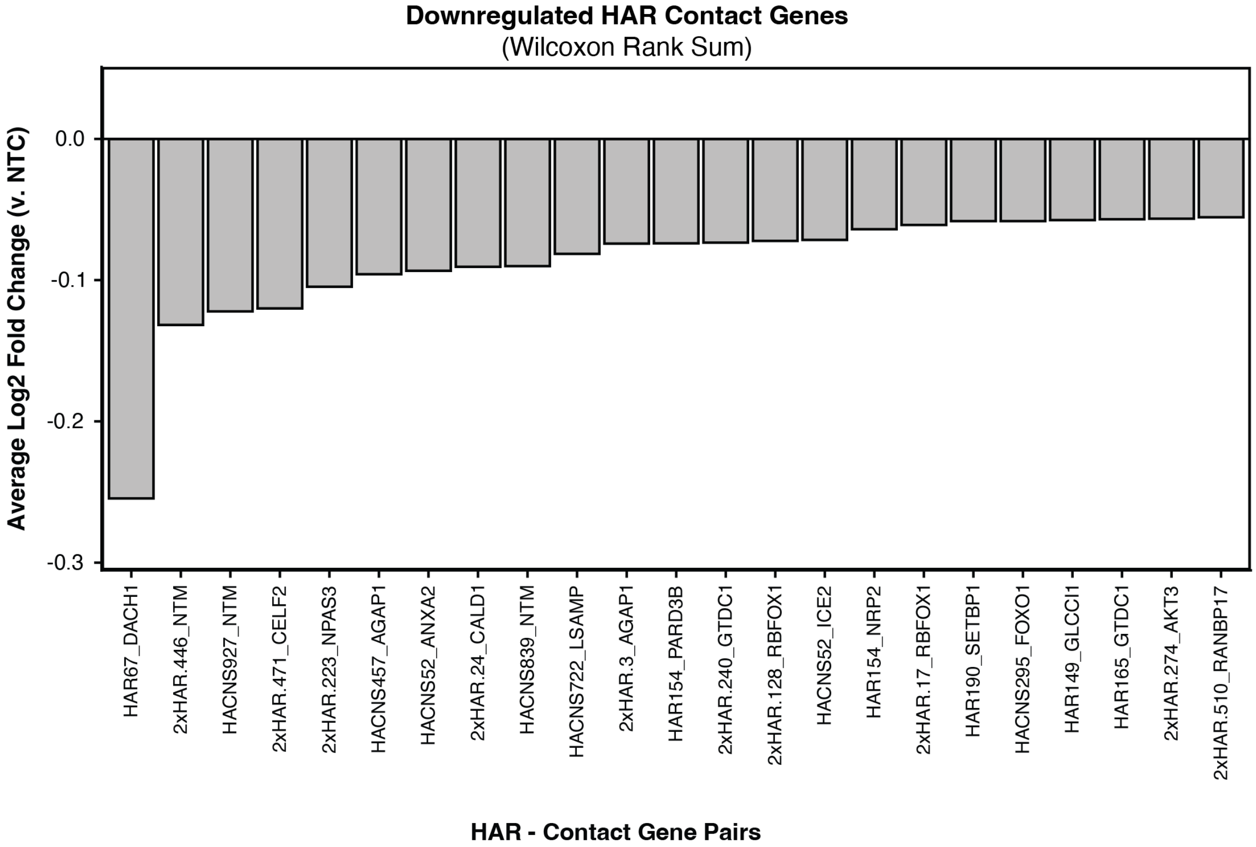


**Fig S5. Differential expression of HAR perturbations and their predicted targets.** Predicted HAR targets were obtained from Capture-C data for HARs in iPSC-derived NSCs (Pal et al. 2024). After differential expression analysis of cells with the same sgRNA target versus NTC guide-bearing cells (Wilcoxon Rank Sum), 23 perturbations (X-axis) showed a log2 fold change > -0.1 (Y-axis) for their predicted targets compared to NTC-guide bearing cells.

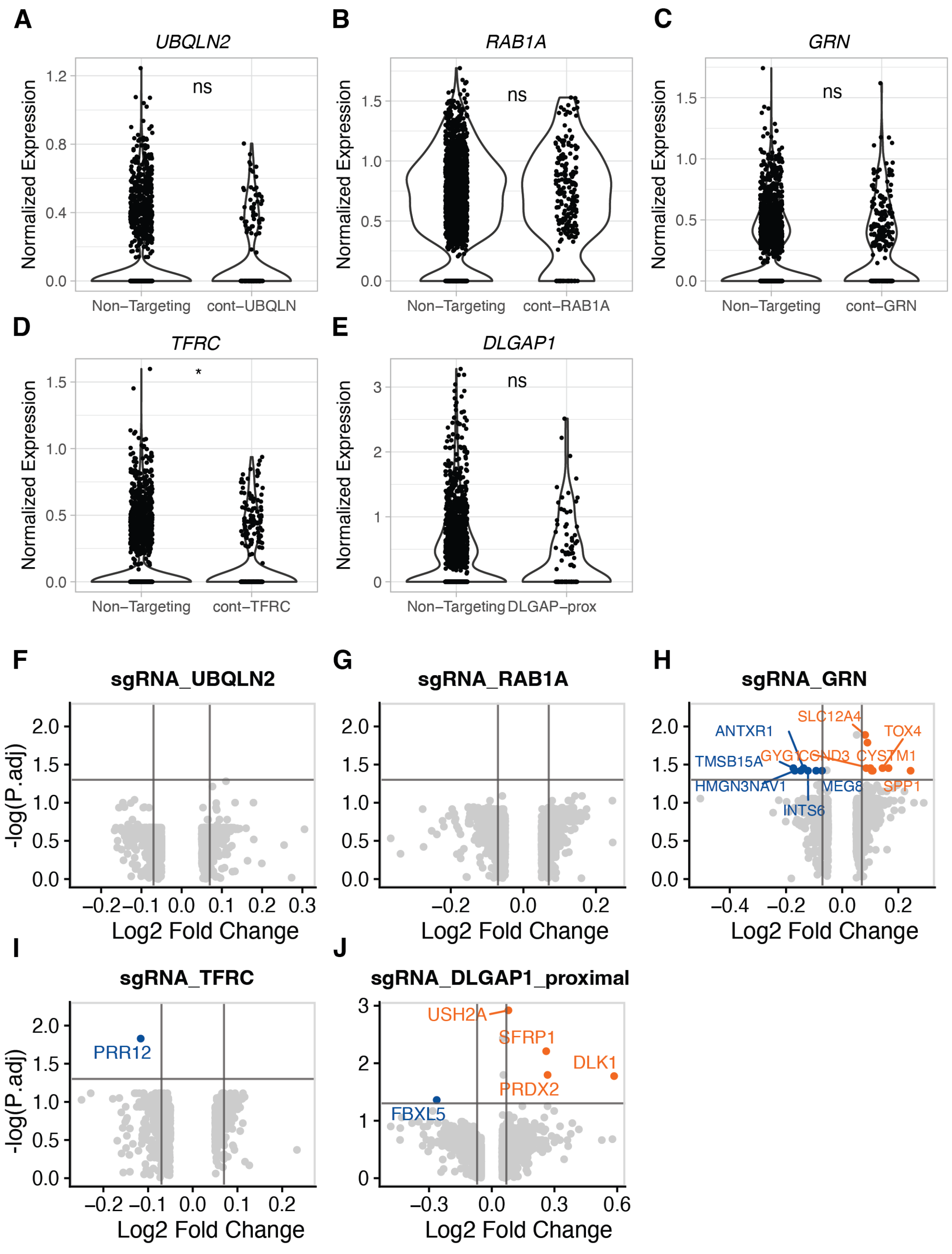


**Fig. S6. Differential expression and power to detect direct gene perturbations in gene-targeting controls. A-E.** Violin plots of target gene expression for guides bearing sgRNAs targeting gene promoters. Nominal significance is shown from Wilcoxon Rank Sum. **F-J**. Volcano plots of differential expression profiles for each of the promoter-targeting sgRNAs, comparing target sgRNA-bearing cells and NTC-sgRNA bearing cells. Adjusted P-values are FDR corrected p-values from Wilcoxon Rank Sum tests, and shown on the Y-axis, and Log2 fold changes are plotted in the X-axis. Significantly downregulated genes are shown in blue, and significantly upregulated genes are shown in orange.

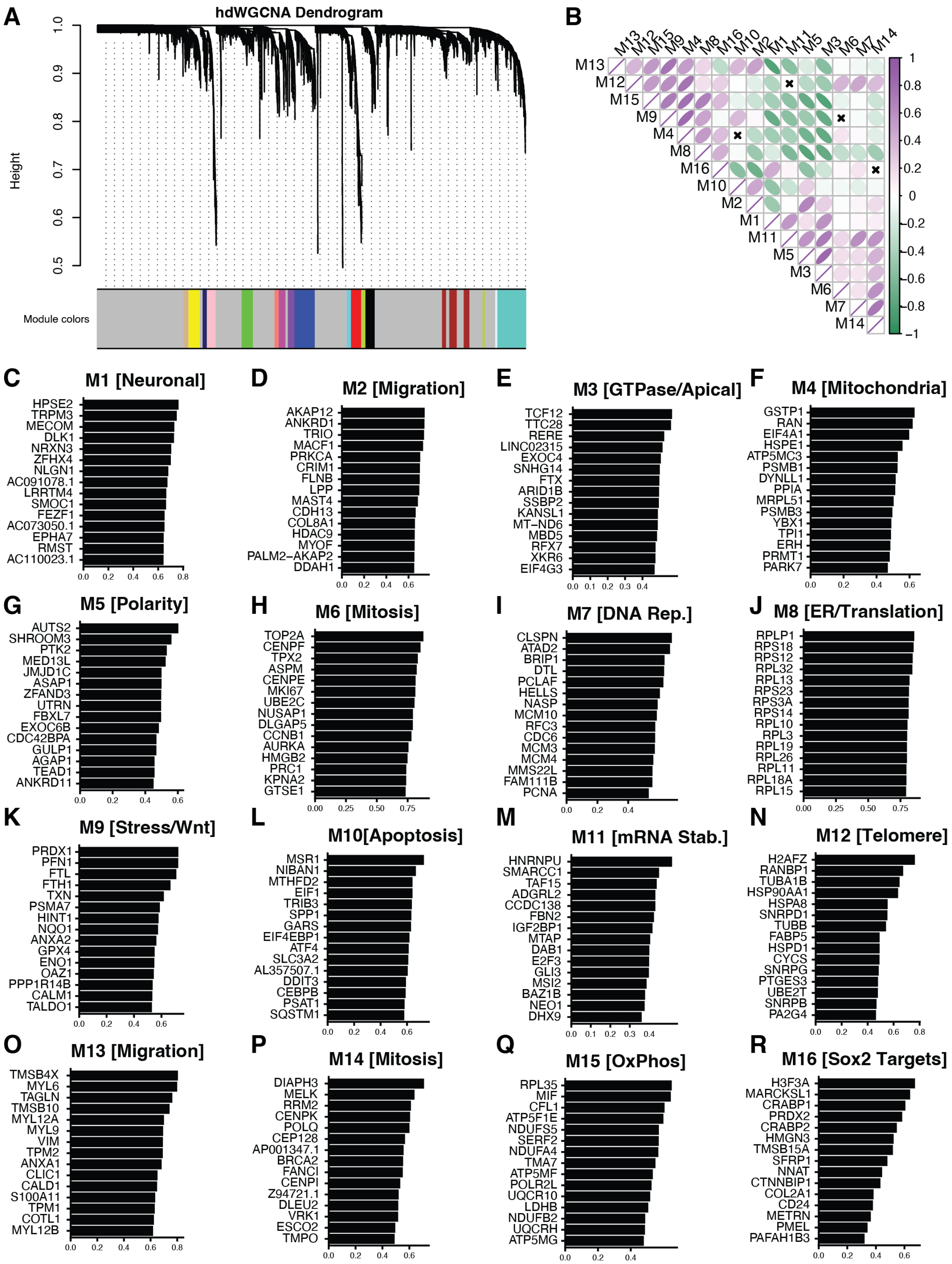

A
hdWGCNA Dendrogram
Height
Module colors
B
M13 M12 M15 M9 M4 M8 M16 M10 M2 M1 M11 M5 M3 M6 M7 M14
C
M1 [Neuronal]
HPSE2
TRPM3
MECOM
DLK1
NRXN3
ZFHX4
NLGN1
AC091078.1
LRRTM4
SMOC1
FEZF1
AC073050.1
EPHA7
RMST
AC110023.1
D
M2 [Migration]
AKAP12
ANKRD1
TRIO
MACF1
PRKCA
CRIM1
FLNB
LPP
MAST4
CDH13
COL8A1
HDAC9
MYOF
PALM2−AKAP2
DDAH1
E
M3 [GTPase/Apical]
TCF12
TTC28
RERE
LINC02315
EXOC4
SNHG14
FTX
ARID1B
SSBP2
KANSL1
MT−ND6
MBD5
RFX7
XKR6
EIF4G3
F
M4 [Mitochondria]
GSTP1
RAN
EIF4A1
HSPE1
ATP5MC3
PSMB1
DYNLL1
PPIA
MRPL51
PSMB3
YBX1
TPI1
ERH
PRMT1
PARK7
G
M5 [Polarity]
AUTS2
SHROOM3
PTK2
MED13L
JMJD1C
ASAP1
ZFAND3
UTRN
FBXL7
EXOC6B
CDC42BPA
GULP1
AGAP1
TEAD1
ANKRD11
H
M6 [Mitosis]
TOP2A
CENPF
TPX2
ASPM
CENPE
MKI67
UBE2C
NUSAP1
DLGAP5
CCNB1
AURKA
HMGB2
PRC1
KPNA2
GTSE1
I
M7 [DNA Rep.]
CLSPN
ATAD2
BRIP1
DTL
PCLAF
HELLS
NASP
MCM10
RFC3
CDC6
MCM3
MCM4
MMS22L
FAM111B
PCNA
J
M8 [ER/Translation]
RPLP1
RPS18
RPS12
RPL32
RPL13
RPS23
RPS3A
RPS14
RPL10
RPL3
RPL19
RPL26
RPL11
RPL18A
RPL15
K
M9 [Stress/Wnt]
PRDX1
PFN1
FTL
FTH1
TXN
PSMA7
HINT1
NQO1
ANXA2
GPX4
ENO1
OAZ1
PPP1R14B
CALM1
TALDO1
L
M10[Apoptosis]
MSR1
NIBAN1
MTHFD2
EIF1
TRIB3
SPP1
GARS
EIF4EBP1
ATF4
SLC3A2
AL357507.1
DDIT3
CEBPB
PSAT1
SQSTM1
M
M11 [mRNA Stab.]
HNRNPU
SMARCC1
TAF15
ADGRL2
CCDC138
FBN2
IGF2BP1
MTAP
DAB1
E2F3
GLI3
MSI2
BAZ1B
NEO1
DHX9
N
M12 [Telomere]
H2AFZ
RANBP1
TUBA1B
HSP90AA1
HSPA8
SNRPD1
TUBB
FABP5
HSPD1
CYCS
SNRPG
PTGES3
UBE2T
SNRPB
PA2G4
O
M13 [Migration]
TMSB4X
MYL6
TAGLN
TMSB10
MYL12A
MYL9
VIM
TPM2
ANXA1
CLIC1
CALD1
S100A11
TPM1
COTL1
MYL12B
P
M14 [Mitosis]
DIAPH3
MELK
RRM2
CENPK
POLQ
CEP128
AP001347.1
BRCA2
FANCI
CENPI
Z94721.1
DLEU2
VRK1
ESCO2
TMPO
Q
M15 [OxPhos]
RPL35
MIF
CFL1
ATP5F1E
NDUFS5
SERF2
NDUFA4
TMA7
ATP5MF
POLR2L
UQCR10
LDHB
NDUFB2
UQCRH
ATP5MG
R
M16 [Sox2 Targets]
H3F3A
MARCKSL1
CRABP1
PRDX2
CRABP2
HMGN3
TMSB15A
SFRP1
NNAT
CTNNBIP1
COL2A1
CD24
METRN
PMEL
PAFAH1B3

**Fig S7. hdWGCNA module composition, expression correlations, and hub genes. A.** Dendrogram of gene correlations within the hdWGCNA modules. **B.** Correlogram showing the correlation of every module's expression (hME) against all other modules. **C-R.** The 15 genes with the highest kME values (plotted on the Y-axes; kME is defined as the correlation of the gene's expression with their module's expression) within each module, i.e. the top 15 hub genes for each module.

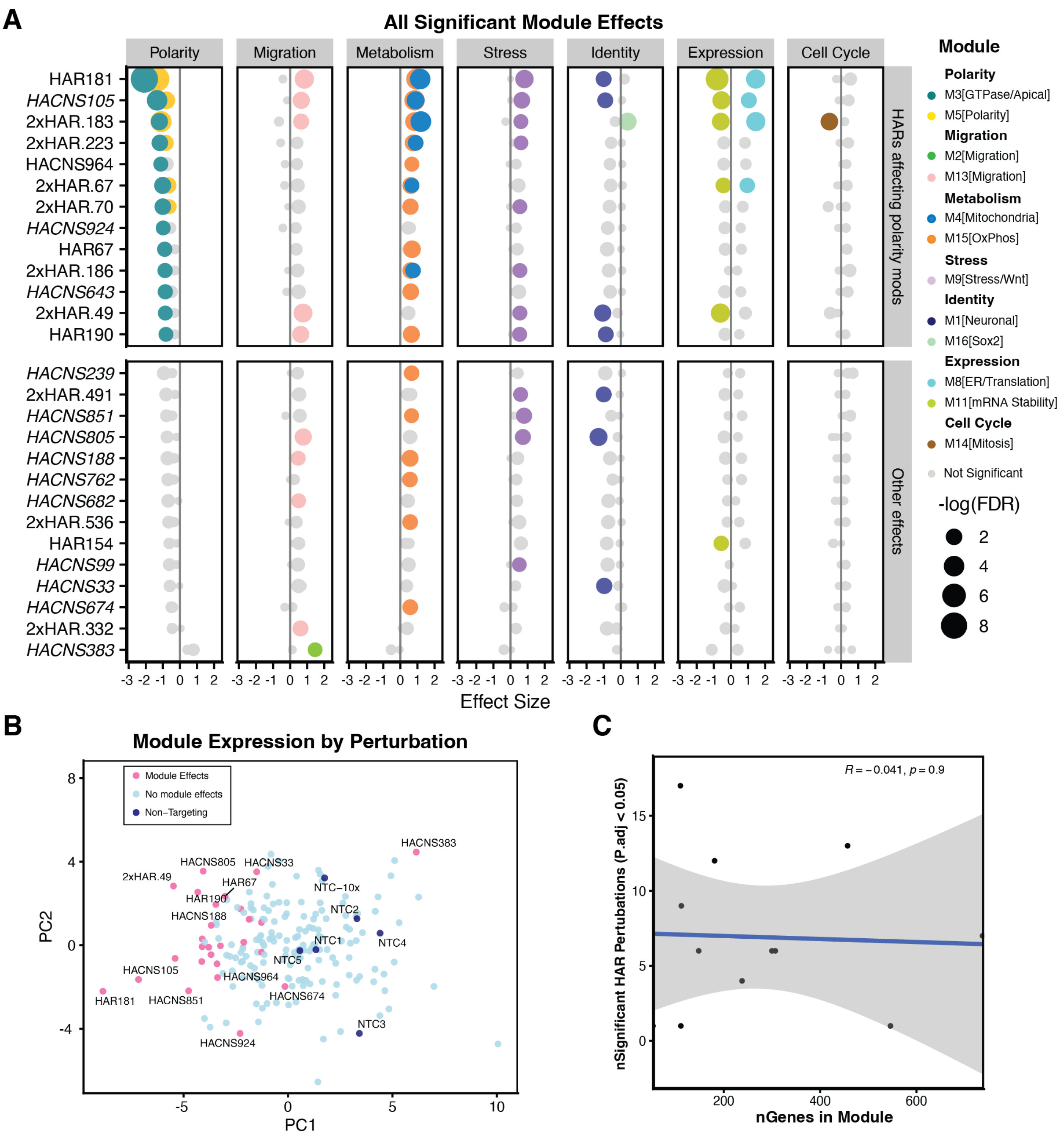


**Fig S8. Multiple linear regression of HAR perturbation effects on modules. A.** Module expression effects of all HAR perturbations which had ≥ 1 significant effect on a module's expression as identified by the MLR model. This plot is an expansion of Figure 3C. Effect sizes are plotted on the X-axis. Significantly affected (FDR < 0.05) modules are colored by module identity as in Figure 3A. **B.** Principal component analysis for the average module eigengene values within each perturbation. Perturbation profiles with significant effects from the MLR are highlighted in pink, whereas module hME profiles for NTC guide-bearing cells are indicated in dark blue. **C.** No significant correlation was found between module size (nGenes) and the number of HAR perturbations resulting in significant effects for that module.

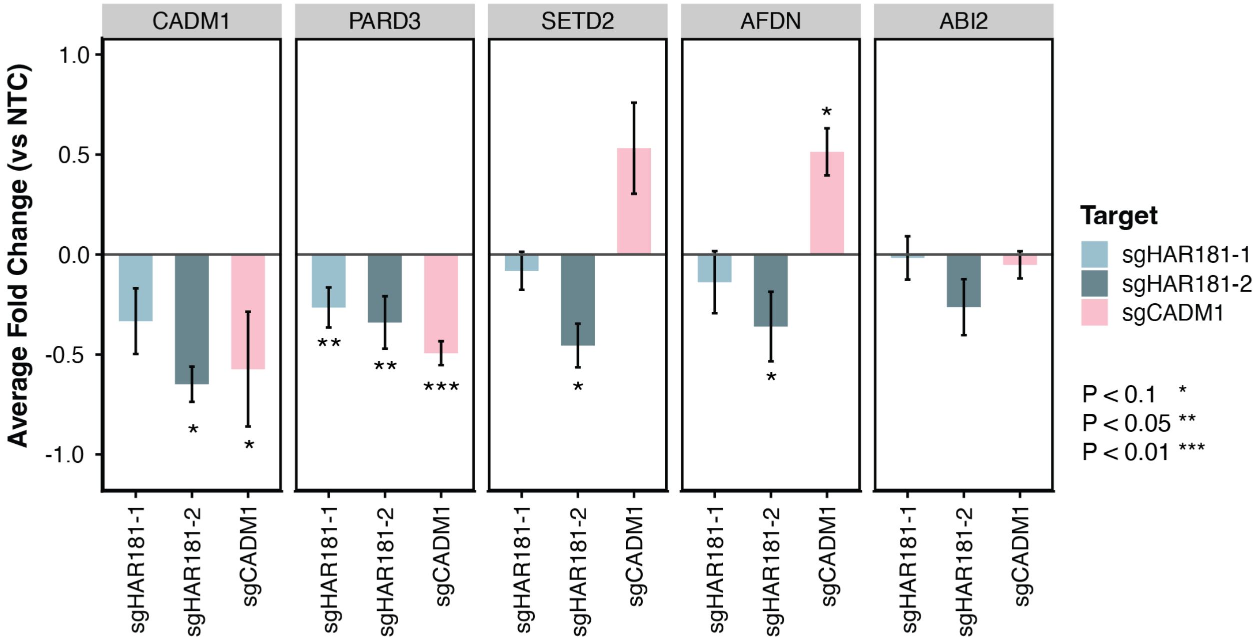


**Fig S9. RT-qPCR of polarity and cytoskeletal genes in an independent cell line.** Average log2 fold change measured by RT-qPCR in bulk dCas9:KRAB CRISPRi assays in an independent human iPSC-derived NSC line. Both guides targeting HAR181 from the original Perturb-seq screen were used, alongside an sgRNA targeting the promoter of the HAR181 target gene, *CADM1*. CT values were normalized to *TBP* expression, then to expression values in cells transfected with an NTC sgRNA. Values are the average fold change of two replicates. P-values were calculated by T-tests comparing NTC values to experimental values.

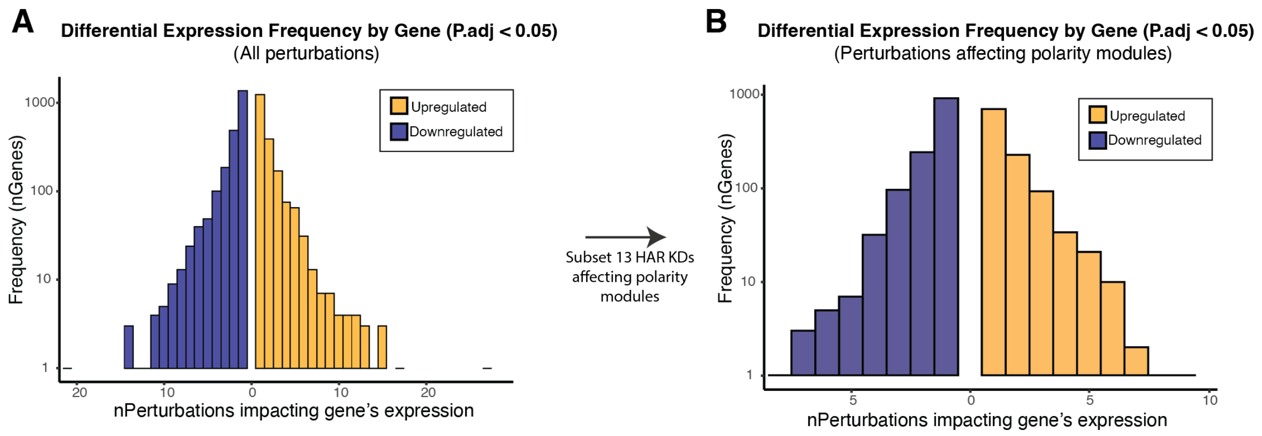


**Fig S10. Distribution of overlapping significant DE genes from 1 or more perturbations.** Histograms of the frequency of the number of different HAR perturbations significantly affecting a given DE gene across the entire dataset (**A**), or across the subset of 13 perturbations which significantly downregulated modules associated with apicobasal polarity (**B**).

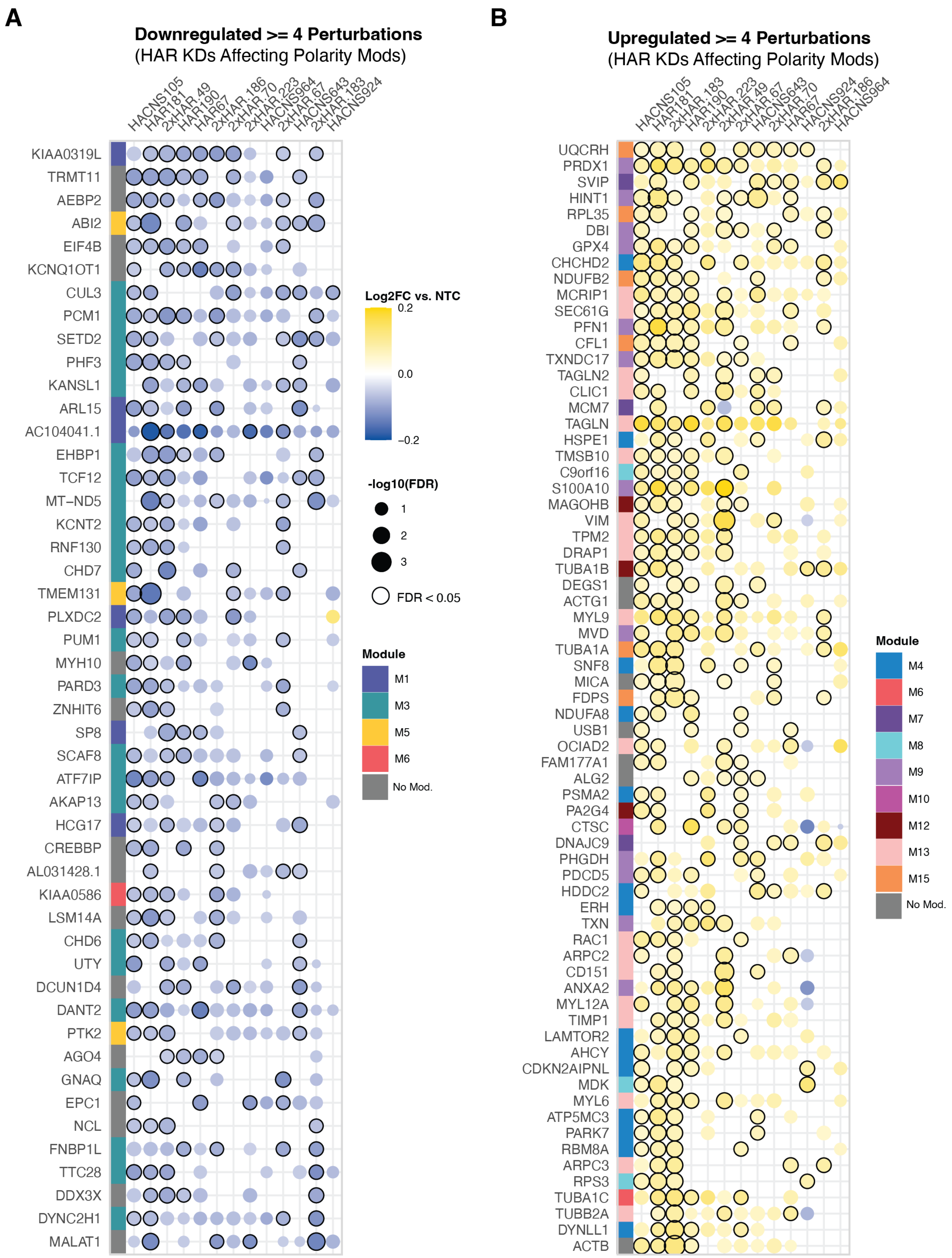

A
Downregulated >= 4 Perturbations
(HAR KDs Affecting Polarity Mods)
HACNS105
HAR181
2xHAR.49
HAR190
HAR67
2xHAR.186
2xHAR.70
2xHAR.223
HACNS964
2xHAR.67
HACNS643
2xHAR.183
HACNS924
KIAA0319L
TRMT11
AEBP2
ABI2
EIF4B
KCNQ1OT1
CUL3
PCM1
SETD2
PHF3
KANSL1
ARL15
AC104041.1
EHBP1
TCF12
MT−ND5
KCNT2
RNF130
CHD7
TMEM131
PLXDC2
PUM1
MYH10
PARD3
ZNHIT6
SP8
SCAF8
ATF7IP
AKAP13
HCG17
CREBBP
AL031428.1
KIAA0586
LSM14A
CHD6
UTY
DCUN1D4
DANT2
PTK2
AGO4
GNAQ
EPC1
NCL
FNBP1L
TTC28
DDX3X
DYNC2H1
MALAT1
Log2FC vs. NTC
0.2
0.0
−0.2
-log10(FDR)
1
2
3
FDR < 0.05
Module
M1
M3
M5
M6
No Mod.
B
Upregulated >= 4 Perturbations
(HAR KDs Affecting Polarity Mods)
HACNS105
HAR181
2xHAR.183
HAR190
2xHAR.223
2xHAR.49
2xHAR.67
HACNS643
2xHAR.70
HAR67
HACNS924
2xHAR.186
HACNS964
UQCRH
PRDX1
SVIP
HINT1
RPL35
DBI
GPX4
CHCHD2
NDUFB2
MCRIP1
SEC61G
PFN1
CFL1
TXNDC17
TAGLN2
CLIC1
MCM7
TAGLN
HSPE1
TMSB10
C9orf16
S100A10
MAGOHB
VIM
TPM2
DRAP1
TUBA1B
DEGS1
ACTG1
MYL9
MVD
TUBA1A
SNF8
MICA
FDPS
NDUFA8
USB1
OCIAD2
FAM177A1
ALG2
PSMA2
PA2G4
CTSC
DNAJC9
PHGDH
PDCD5
HDDC2
ERH
TXN
RAC1
ARPC2
CD151
ANXA2
MYL12A
TIMP1
LAMTOR2
AHCY
CDKN2AIPNL
MDK
MYL6
ATP5MC3
PARK7
RBM8A
ARPC3
RPS3
TUBA1C
TUBB2A
DYNLL1
ACTB
Module
M4
M6
M7
M8
M9
M10
M12
M13
M15
No Mod.

**Fig S11. Convergence in DE genes among ≥ 4 perturbations affecting polarity modules**. Plots showing the all DEGs affected by at least 4 HAR perturbations in the subset of 13 perturbations which downregulated polarity-associated modules. These are an extension of Figure 5C-D. Among perturbations with negative (**A**) or positive (**B**) effect sizes on polarity-associated modules M3 and M5, the DEGs are shown with Log2 fold change indicated by circle color, -log(FDR) indicated by circle size, and significant (FDR < 0.05) values outlined in black. Each gene's module membership is indicated by the color in the left ribbon. Genes are arranged by the frequency with which they are affected by HAR perturbations in this subset.

A

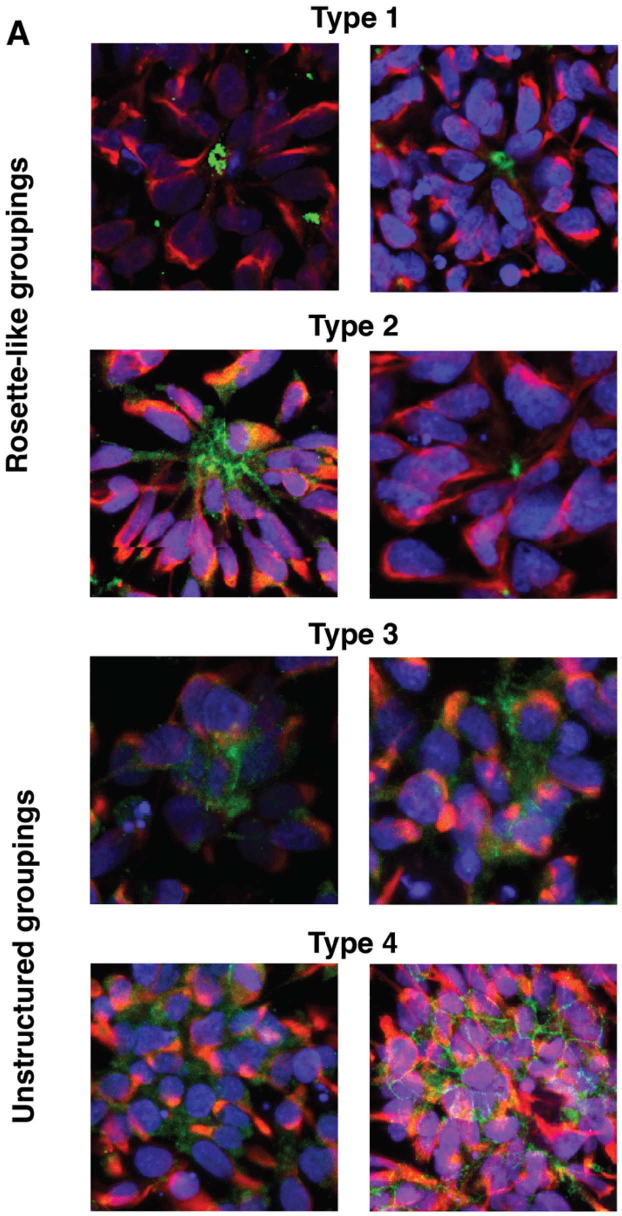


**Type 1**

- Cells are radially oriented around a single maxima of green (PARD3) signal.

**Type 2**

- Cells are primarily oriented around a single maxima of green, with some aberrations.
- Central green signal (PARD3) may be abornally shaped, but is not distributed throughout grouping.

**Type 3**

- Cells are grouped around a single centroid, but are not oriented radially.
- Green signal (PARD3) may be distributed around mass.

**Type 4**

- Cells are grouped with no centroid, forming a mass of cells without distinct orientation.
- Some cells in this mass are entirely engrailed by green signal (PARD3).
- Green signal is distributed throughout mass.

B

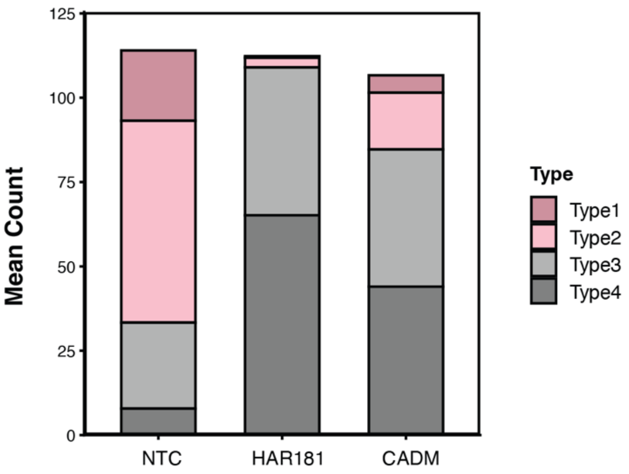

**Fig S12. Blinded scoring and quantification of NSC grouping morphologies. A.** Rubric for scorers to determine cell grouping types. Groupings were defined as 4 or more adjoined cells marked by PARD3. Types 1 and 2 represent structured rosette-like cell grouping morphologies. Types 3 and 4 are unstructured or disordered grouping morphologies. **B.** Bar chart showing the mean counts of each cell-grouping type in each condition across 3 evaluators and 2 replicates. Grouping morphology types are indicated by bar color.
